# Recent thymic emigrants produce antimicrobial IL-8, express complement receptors and are precursors of a tissue-homing Th8 lineage of memory cells

**DOI:** 10.1101/059535

**Authors:** Marcin L Pekalski, Arcadio Rubio García, Ricardo C Ferreira, Daniel B Rainbow, Deborah J Smyth, Meghavi Mashar, Jane Brady, Natalia Savinykh, Xaquin Castro Dopico, Sumiyya Mahmood, Simon Duley, Helen E Stevens, Neil M Walker, Antony J Cutler, Frank Waldron-Lynch, David B Dunger, Claire Shannon-Lowe, Alasdair J Coles, Joanne L Jones, Chris Wallace, John A Todd, Linda S Wicker

## Abstract

The adaptive immune system utilizes multiple mechanisms linked to innate immune cell functions to respond appropriately to pathogens and commensals. Here we discover further aspects of this connectivity by demonstrating that naïve T cells as they emerge from the thymus (recent thymic emigrants, RTEs) express complement receptors (CR1 and CR2), the bacterial pathogen recognition receptor TLR1 and an enzyme that deactivates bacterial lipopolysaccharide (AOAH) and following activation secrete the anti-microbial cytokine IL-8. CR2^+^ naïve T cells also express a selection of genes associated with tissue migration, consistent with the hypothesis that following emigration from the thymus RTEs seed peripheral compartments where some pursue their anti-microbial potential by becoming IL-8-producing CR2^+^ memory cells while others undergo homeostatic expansion. CR2^+^ naïve and memory cells are abundant in children but decrease with age, coinciding with the involution of the thymus. The ability of CR2, which is also a receptor for Epstein-Barr Virus (EBV), to identify recent thymic emigrants will facilitate assessment of thymic function during aging and aid investigations of multiple clinical areas including the occurrence of T cell lymphomas caused by EBV.

## INTRODUCTION

The maintenance of a diverse, naïve T cell repertoire arising from the thymus (recent thymic emigrants, RTEs) is critical for health (1). Thymic involution decreases naïve CD4^+^ T cell production with age in humans and is compensated for by the homeostatic expansion of naïve cells that have emigrated from the thymus earlier in life (2, 3) and seed peripheral compartments (4). Naïve T cells that have undergone decades of homeostatic expansion show reduced T cell receptor diversity, which has the potential to negatively impact host (1, 5). CD31 (PECAM-1) expression identifies cells that have divided more often in the periphery (CD31^−^) from those that have not (CD31^+^), although CD31^+^ T cells still divide with age as evidenced by the dilution of signal joint T-cell receptor rearrangement excision circles (sjTRECs) (6). The tyrosine-protein kinase-like 7 receptor (encoded by *PTK7*) has been reported as a marker of RTEs within the CD31^+^ naïve T cell subset. However, PTK7 expression reduces on naïve CD4^+^ T cells with donor age (7, 8) thereby limiting its usefulness to purify the most naïve T cells present in adults for further study. We and others have reported (9, 10) that CD25 is upregulated in naïve T cells that have expanded in the periphery and that the proportion of naïve CD4^+^ T cells that are CD31^+^ CD25^+^ or CD31^−^ CD25^+^ increases with age. CD31^+^ CD25^−^ naïve CD4^+^ T cells contain the highest content of sjTRECs as compared to their CD31^−^ and CD25^+^ counterparts thereby identifying the CD31^+^ CD25^−^ naïve CD4^+^ T cell subset as containing the highest proportion of RTEs in adults (9).

In the current study we isolated four naïve CD4^+^ T cell subsets from 20 adults based on CD31 and CD25 expression and conducted transcriptome profiling to determine their molecular signatures. These signatures define gene expression patterns in naïve T cells as they first undergo post-thymic maturation and then continue to expand over years and decades in the periphery waiting to encounter their cognate antigen. Owing to this systematic approach, we made an unexpected discovery that complement receptor 2 (CR2) in conjunction with CD31 and CD25 defines the most naïve CD4^+^ T cells within adults and that CR2 is strongly expressed on naïve T cells in neonates and children. In patients treated with alemtuzumab we observed that during homeostatic reconstitution, newly emerging naïve T cells from the adult thymus express high levels of CR2 and ∼50% produce IL-8, phenotypes that are comparable to those of naïve T cells in neonates. Therefore, analysis of CR2 levels on naïve CD4^+^ T cells and CD8+ T cells should provide a new tool in assessing thymic function as people age and during bone marrow transplantation (11), HIV infection (12) and immune reconstitution following immune depletion (13) or chemotherapy (14). Recently a role for other complement receptors in T cell immunity has been described (15). In contrast to the regulatory function of CR1 and CR2, the C3a and C5a receptors in T cells stimulate inflammasome activity and effector functions (16). Our results also support the existence of an IL-8-producing memory T cell subset identified in a recent study (17), which may, dependent on future investigations, prove to be a Th8 lineage. Based on the transcription signature shared by CR2^+^ RTEs and CR2^+^ memory cells that includes the expression of transcription factor ZNF462 and genes associated with tissue migration, we suggest that CR2^+^ RTEs surveying peripheral tissues are precursors of a Th8-like subset of memory cells.

## RESULTS

### Naïve CD4^+^ T-cell Subsets Defined by CD31 and CD25 Differentially Express *Complement Receptor 2 (CR2)* and *AOAH*, encoding the enzyme acyloxyacyl hydrolase that inactivates LPS

We assessed the proportion of four naïve CD4^+^ T cell subsets defined by CD31 and CD25 expression (9) in neonates, children and adults in a population study of 391 donors (**Figure 1A,B**). CD31^+^ CD25^−^ naïve CD4^+^ T cells decreased with age and this decrease was compensated for by the homeostatic expansion of three subsets of naïve T cells: CD31^+^ CD25^+^, CD31^−^ CD25^−^ and CD31^−^ CD25^+^. As expected (18), the proportion of both naïve CD4^+^ and CD8^+^ T cells negatively correlated with donor age (**Figure S1A**). To define molecules associated with the least expanded naïve subset, we performed a statistically powered, genome-wide RNA analysis of FACS-purified naïve CD4^+^ T cells from 20 adults sorted into four subsets based on CD31 and CD25 expression (**Figure S1B**). Principal component analysis of differentially expressed genes amongst the four subsets showed a clear separation between the groups, particularly between CD31^+^ cells and their CD31^−^ counterparts (**Figure S1C**). Genes with higher expression in CD31^+^ CD25^−^ naïve cells as compared to the CD31^−^ CD25^−^ subset included two genes not normally associated with T cells: *AOAH*, which encodes the enzyme acyloxyacyl hydrolase that inactivates LPS and is highly expressed in innate immune cells (19), and *CR2*, which encodes a cell surface protein that binds C3d and other complement components (20) and is also a receptor for EBV in humans (21) (**Figure 1C, Spreadsheets S1A-D**). The expression of 12 genes, including *CR2, AOAH, TOX* (a transcription factor reported to regulate T-cell development in the thymus (22)) and *CACHD1*, an uncharacterized gene that may encode a protein that regulates voltage-dependent calcium channels, was lower when CD31^+^ CD25^−^ cells either lost expression of CD31 (**Figure 1C**) or up-regulated CD25 (**Figure S1D**, see list of shared up- or down-regulated genes upon expansion in **Spreadsheet S1E**).

**Figure 1.**
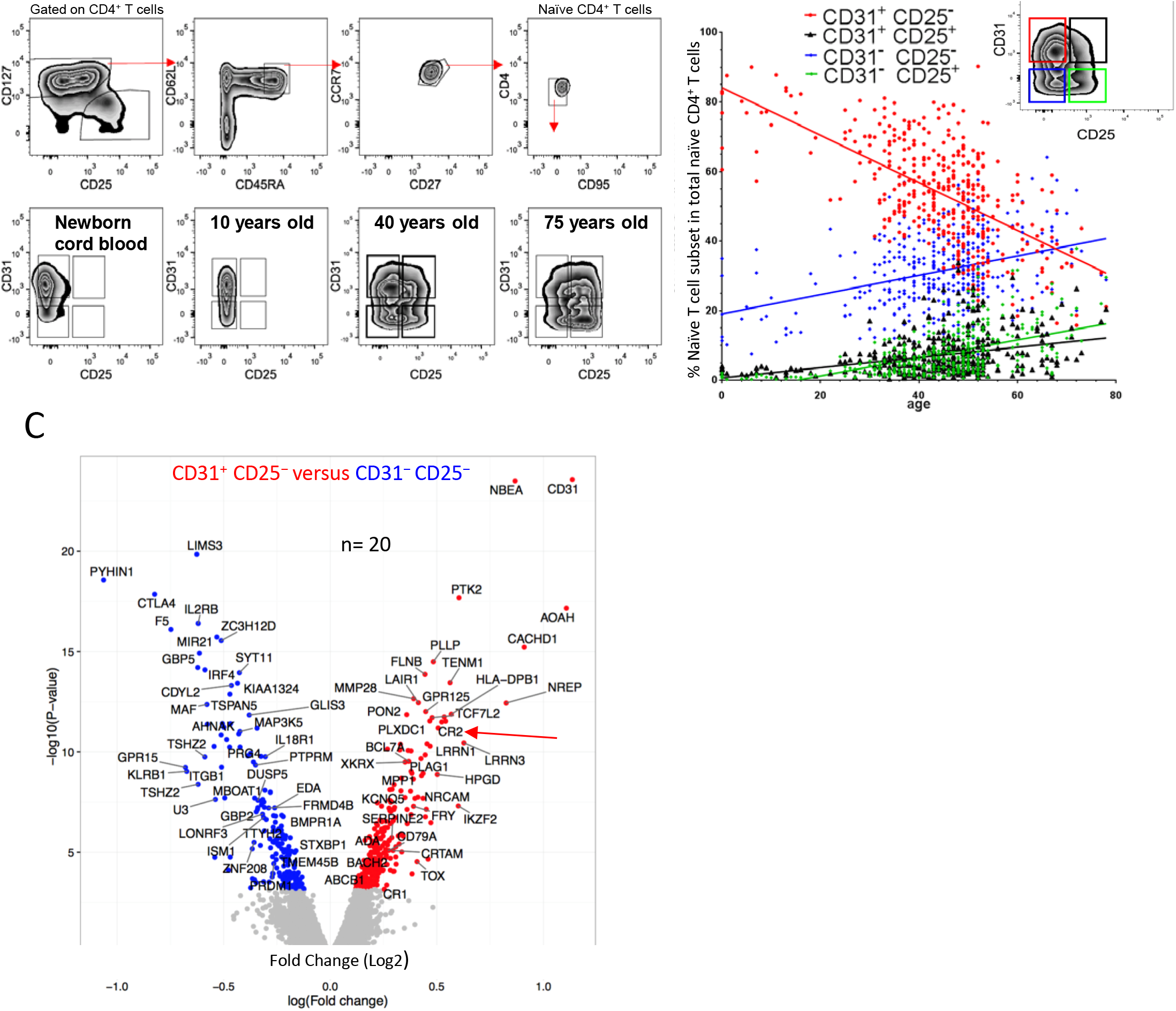
Gene expression profiling of four naïve CD4^+^ T cell subsets identify age-related molecular signatures. **(A)** Gating strategy defining human naïve CD4^+^ T cells; naïve T cells were further stratified by CD31 and CD25. Representative examples (from N=391) of naïve CD4^+^ T cells. **(B)** The proportion of naïve CD4^+^ T cells as a function of age (color coding shown above graph). **(C)** Volcano plot of differences in gene expression (microarray platform) between CD31^+^ CD25^−^ and CD31^−^ CD25^−^ naïve CD4^+^ T cells; red and blue symbols for genes with increased and decreased, respectively, expression in CD31^+^ CD25^−^ naïve CD4^+^ T cells.

Genes up-regulated in expanded CD31^−^ CD25^−^ cells as compared to CD31^+^ CD25^−^ cells (**Figure 1C**). are consistent with the occurrence of activation and differentiation events during the homeostatic proliferation of naïve T cells and include *PYHIN1*, a gene encoding an interferon-induced intracellular DNA receptor that is a member of a family of proteins that induces inflammasome assembly (23, 24), *GBP5*, encoding a protein that promotes NLRP3 inflammasome assembly (25), *CTLA4, KLRB1* (encoding CD161) a marker of IL-17 T cells (26) and *IL2RB*, as well as transcription factors such as *IRF4, PRDM1* (encoding BLIMP-1) and *MAF*.

### Naïve T Cells Expressing CR2 are More Prevalent in Children than Adults

We verified the microarray results using flow cytometric analysis, with the CD31^+^ CD25^−^ naïve CD4^+^ T cell subset having the highest proportion of cells positive for CR2 (**Figure 2A,B**). The three subsets previously shown to be products of homeostatic expansion based on sjTREC content (9) had lower proportions of CR2^+^ cells. The reduction in the percentage of CR2^+^ naïve cells within the CD31^+^ CD25^−^ subset by age (**Figure 2B**) was more pronounced when considered out of total CD4^+^ T cells since the proportion of naïve cells within the CD4 population decreases with age (**Figure S2A**). The CR2^+^ fraction of the CD31^+^ CD25^−^ naïve CD4^+^ T cell subset has the highest level of CR2 on a per cell basis and CR2 density on this subset decreases with age (**Figure 2A**). A similar pattern of a decreasing proportion of CR2^+^ naïve cells and reduced density of CR2 per cell with age was also observed on naïve CD8^+^ T cells (**Figure S2B**). Because PTK7 has been described as a marker of RTE (7, 8), we examined our microarray data for differential expression in the four subsets of naïve cells in adults to determine if a similar pattern to that observed for *CR2* could be observed. Although no differential expression was evident in any of the comparisons (**Table S1**), this result could be due to the low level of PTK7 expression in adult cells (see Figure 2 in (7)) together with the sensitivity limits of the microarray platform. We confirmed the low expression of PTK7 on the surface of adult naïve CD4^+^ T cells using cord blood cells as a positive control for PTK7 staining (**Figure S2C,D**).

**Figure 2.**
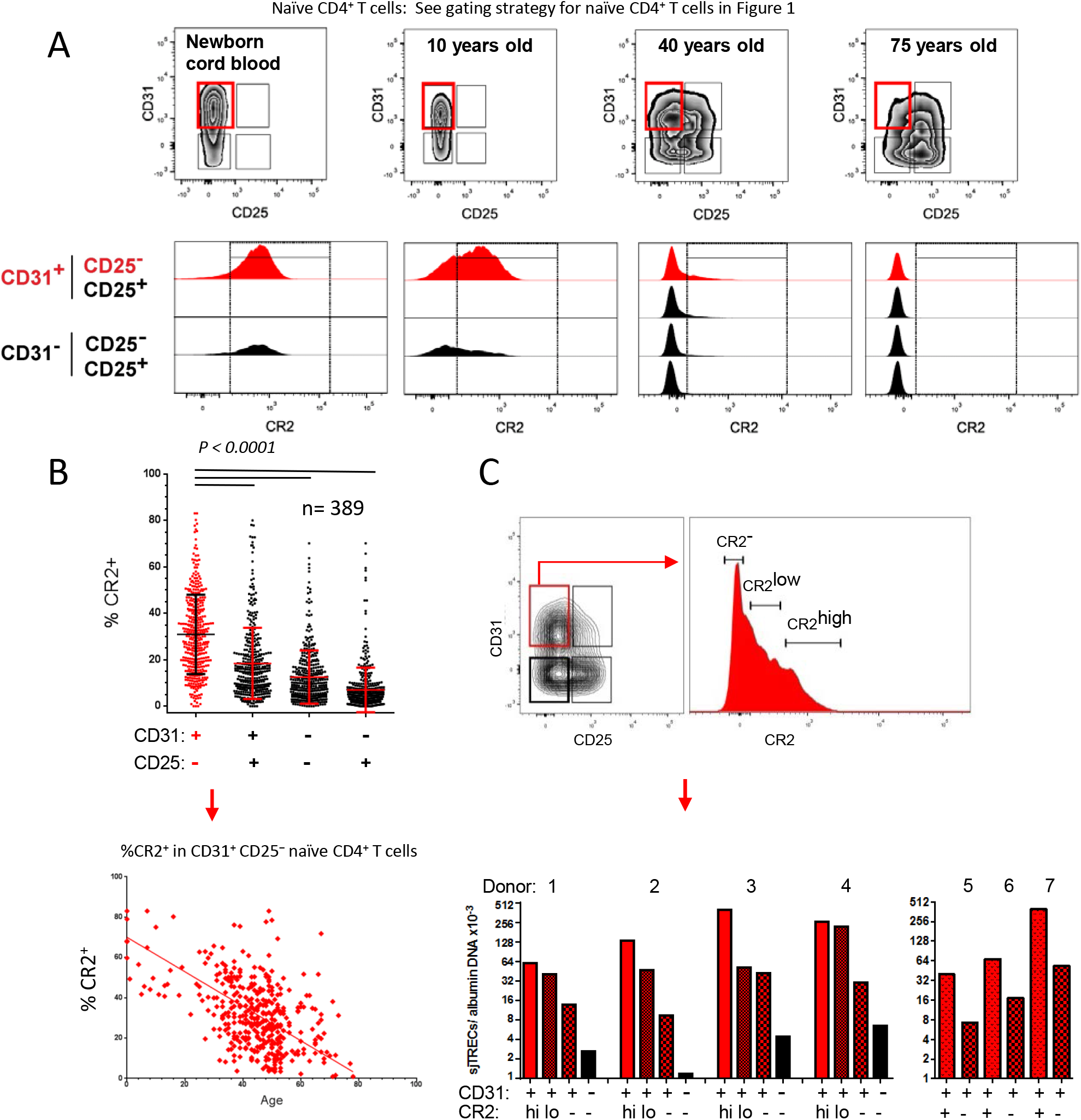
CR2 marks the most naïve CD4^+^ T cell subset. **(A)** Representative examples (from N=389) of CR2 expression in naïve T cell subsets. **(B)** %CR2^+^ cells in each subset and frequency of CR2^+^ cells in the CD31^+^ CD25^−^ naïve CD4^+^ T cell subset as a function of age. **(C)** Representative sorting strategy for CD31^+^ CD25^−^ naïve CD4^+^ T cells identified as CR2^−^, CR2^low^ and CR2^high^ (donors 1-4, for donors 5-7 the CR2^+^ gate is a combination of low and high CR2-expressing cells) that were assessed for sjTRECs.

### CR2^+^ Naïve CD4^+^ T Cells Have a Higher sjTREC Content than Their CR2^−^ Counterparts

To determine whether CR2 is a molecular marker of the subset of CD31^+^ CD25^−^ naïve CD4^+^ T cells that have proliferated the least in the periphery since emigrating from the thymus, we sorted CR2^hl^, CR2^low^ and CR2^−^ CD31^+^ CD25^−^ naïve CD4^+^ T cells along with CD31^−^ CD25^−^ naïve CD4^+^ T cells from four adult donors and, when cell numbers were limiting, CR2^+^ and CR2^−^ CD31^+^ CD25^−^ naïve CD4^+^ T cells from three additional adult donors, and assessed sjTREC levels (**Figure 2C**). As expected based on previous studies showing that the loss of CD31 expression from naïve cells is associated with a substantial loss of sjTRECs (2, 6, 9), sorted CD31^−^ naïve CD4^+^ T cells had fewer sjTRECs than any of the CD31^+^ populations (**Figure 2C**). CR2^+^ cells had more sjTRECs than CR2^−^ cells in all cases (N=7, *P* = 0.0023 and *P* = 0.018 using CR2^hi^ and CR2^lo^, respectively, compared with CR2^−^ CD31^+^ cells for donors 1-4 and CR2^+^ CD31^+^ versus CR2^−^ CD31^+^ cells for donors 5-7, Mann Whitney rank test) demonstrating that in adults, CR2^+^ cells have undergone fewer rounds of homeostatic expansion as compared to CR2^−^ cells within the CD31^+^ CD25^−^ subset of naïve CD4^+^ cells. Where cell numbers were sufficient to separate the CR2^hi^ and CR2^low^ naïve CD4^+^ T cells, CR2^low^ cells had fewer sjTRECs than the CR2^hi^ cells in each comparison (N=4, *P* = 0.11) suggesting that higher CR2 expression on a per cell basis on CD31^+^ CD25^−^ naïve CD4^+^ T cells identifies cells that have divided the least number of times since leaving the thymus. These observations along with our previous demonstration of CD25 on homeostatically expanded naïve T cells (9) explain why naïve CD4^+^ T cells isolated only by CD31 expression show an age-dependent loss of sjTRECs (2, 6).

### RTEs from the Adult Thymus Are CR2^+^

To further test the hypothesis that CR2 expression on naïve T cells defines RTEs throughout life rather than being specific to cells generated during the neonatal period, we monitored newly generated naïve CD4^+^ T cells in eight multiple sclerosis patients depleted of T and B lymphocytes using a monoclonal antibody specific for CD52 (alemtuzumab) (13) (**Figures 3A**). Twelve months after depletion, all eight patients had more naïve CD4^+^ (**Figure 3B**) and CD8_+_ (**Figure S3B**) T cells expressing CR2 as compared to baseline demonstrating that *de novo* RTEs produced in the adult thymus are also defined by CR2 expression. Interim time points were available from most patients (**Figures 3C, S3C**) showing that when the first few naïve CD4^+^ T cells were detected after depletion (3-9 months post-treatment), they were essentially all CR2^+^ with a density of CR2 per cell equalling that seen in cord blood (**Figure 2A**). This observation was independent of whether a patient had good or poor naïve CD4^+^ T cell reconstitution overall. We also observed PTK7 on the surface of naïve CD4^+^ T cells in MS patients reconstituting their T and B cells (**Figure S2E**); PTK7 levels were higher than those observed on naïve cells from healthy adults.

**Figure 3.**
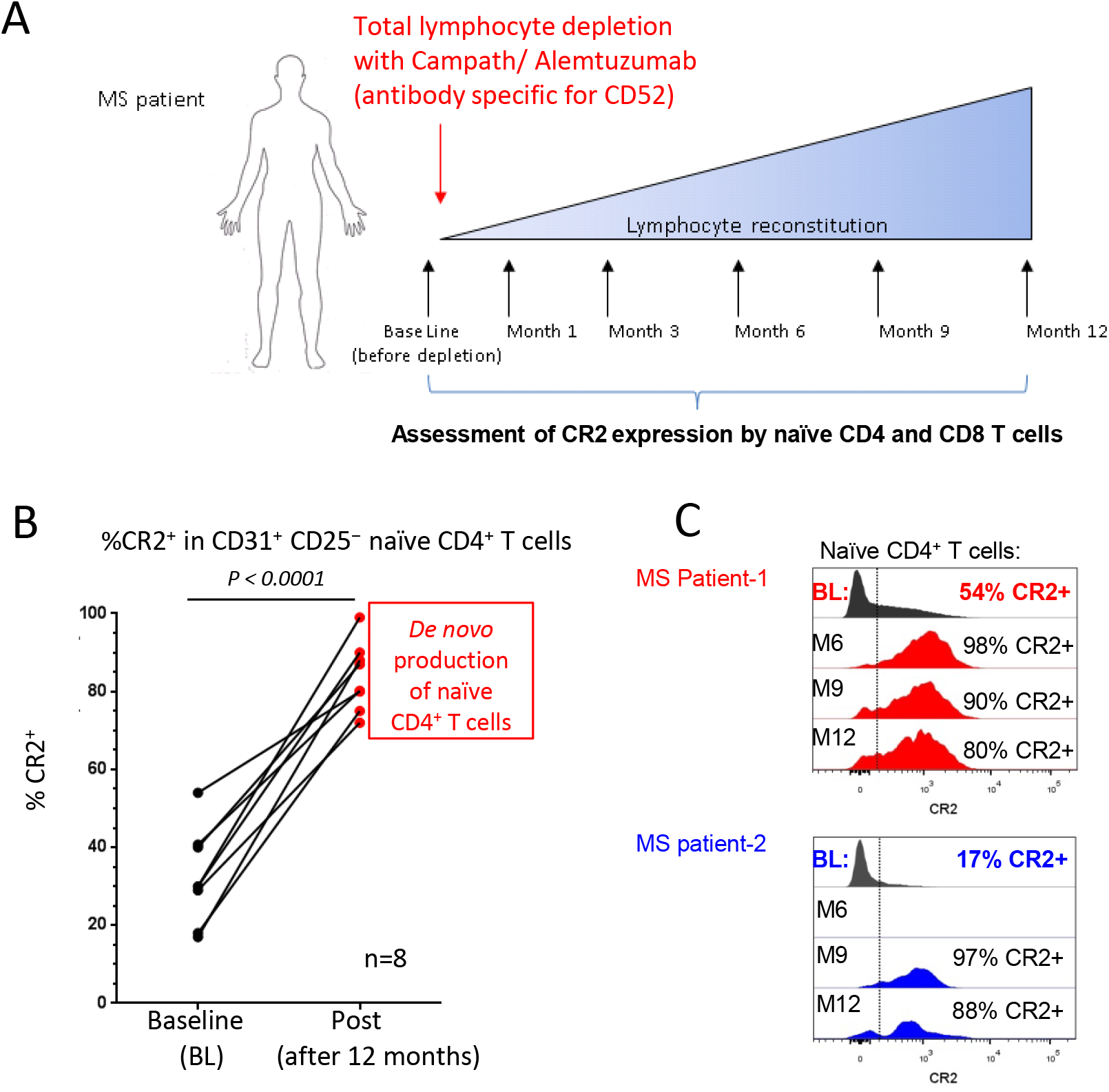
Increased complement receptor 2 (CR2) expression by human naïve CD4^+^ T cells during *de novo* reconstitution. **(A)** Treatment and sampling time points of MS patients. **(B)** Frequency of CD31^+^ CD25^−^ naïve CD4^+^ T cells expressing CR2 in MS patients before (baseline, BL) and 12 months after lymphocyte depletion with anti-CD52, (Campath). **(C)** CR2 expression on CD31^+^ CD25^−^ naïve CD4^+^ T cells from two patients before and at various times during reconstitution; time courses from six additional patients are shown in **Figure S3C**.

A potential utility of our observation is to use CR2 as a biomarker of the functionality of the human thymus. Along these lines we compared the frequency of CR2^+^ cells within the CD31^+^ CD25^−^ naïve CD4^+^ T cell subset prior to lymphocyte depletion with the ability of the thymus to reconstitute the naïve CD4^+^ T cell compartment (**Figure S3A**). The two patients (P2 and P6) who failed to reconstitute their naïve T cell pool to pre-treatment levels by 12 months had the lowest levels of CR2^+^ T cells within the CD31^+^ CD25^−^ naïve CD4^+^ T cell subset (18% and 17%) prior to treatment. In contrast patients who reconstituted their naïve CD4^+^ T cell pool to or above baseline by 12 months had on average 38% (range 29-54%) CR2^+^ cells in the CD31^+^ CD25^−^ naïve CD4^+^ T cell subset prior to treatment. These data are consistent with the hypothesis that adults who have a limited number of CR2-expressing cells in their CD31^+^ CD25^−^ naïve CD4^+^ T cell subset have reduced thymic function and poorly reconstitute their naïve cell pool following immune depletion with anti-CD52.

### Gene Expression Profiling of CR2^+^ and CR2^−^ Naïve Cells Reveals an Innate Immunity Gene Signature

To evaluate the potential function of CR2^+^ naïve CD4^+^ T cells, we compared RNA isolated from sorted CR2^+^ and CR2^−^ CD31^+^ CD25^−^ naïve CD4^+^ T cells *ex vivo* and after activation (**Figure 4A, Spreadsheets S2,3, Tables S1,2**). This strategy revealed a unique transcriptional signature of the CR2^+^ RTEs. *Complement receptor 1 (CR1), AOAH* and *TLR1* were more highly expressed in CR2+ cells. Co-expression of CR1 and CR2 was observed in CD31^+^ CD25^−^ naïve CD4^+^ T cells from healthy controls (**Figure 4B**) and MS patients reconstituting their T cell repertoire post-alemtuzumab treatment (**Figure S4A**). Co-expression of CR1 and CR2 was also observed on naïve CD8^+^ T cells (**Figures S4B,C**). Consistent with CR2 marking RTEs, we observed that *PTK7* mRNA was more abundant in the CR2^+^ as compared to the CR2^−^ CD31^+^ CD25^−^ naïve CD4^+^ T cells although over 20-fold fewer *PTK7* reads were observed as compared to *CR2* mRNA (**Figure S2F**).

**Figure 4.**
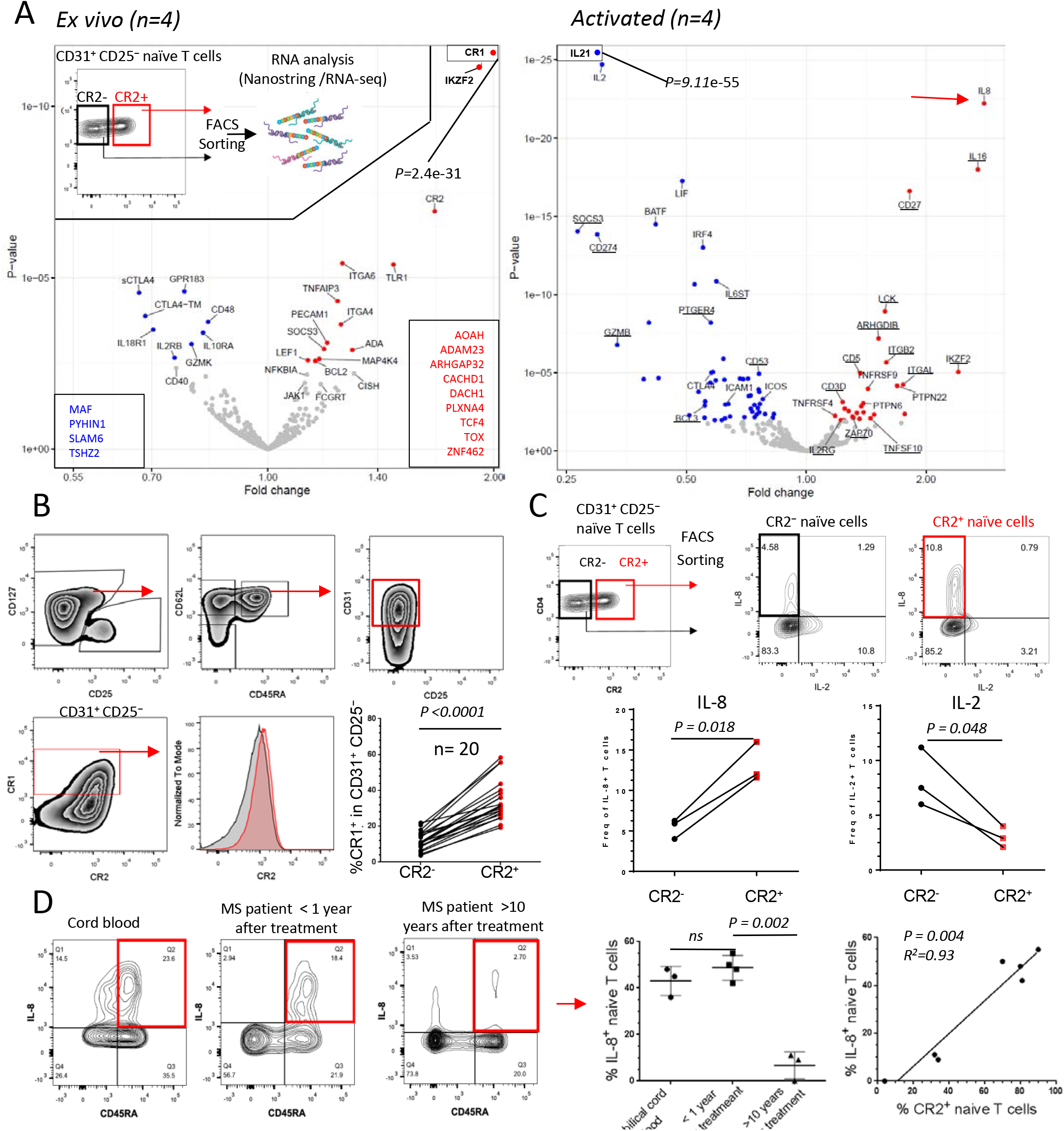
CR2^+^ naïve CD4^+^ T cells have a unique molecular signature. **(A)** Volcano plots of differences in gene expression (NanoString platform) between CR2^+^ versus CR2" naïve CD4^+^ T cells (gating strategy shown as insert) *ex vivo* and after PMA/ionomycin activation. Underlined genes have reduced expression after activation. Genes in boxes are from the RNA-seq platform. **(B)** *Ex vivo* CR1 protein expression on CR2^+^ and CR2" cells using the gating strategy shown (n=20, age range 0 to 17). Representative histograms and compiled frequencies of cytokine production following activation of CR2^+^ and CR2" cells sorted from CD31^+^ CD25" naïve CD4^+^ T cells (n=3, age range 30-45) **(C)** and of naïve (CD45RA^+^ CD31^+^ CD25") T cells in cord blood and adult MS patients (unpaired t test). Correlation of %IL-8-producing and %CR2^+^ CD31^+^ CD25" naïve CD4^+^ T cells in MS patients (n=7) **(D)**.

Following activation, *IL8* mRNA was more highly expressed in CR2^+^ naïve CD4^+^ T cells whereas *IL2, IL21, LIF* and *IFNG* were more highly expressed in CR2^−^ cells (**Figure 4A, Spreadsheet S3, Table S2**), which is consistent with studies showing that RTEs from children secrete less IFN-γ and IL-2 (7) and more IL-8 (8). *TNF, LTA* and *IL23A* were highly upregulated with activation but there was no difference between the CR2**+** and CR2^−^ naïve subsets. *IL8* upregulation was of particular interest since it has been termed a phenotype of neonatal naïve cells (27). We therefore measured IL-8 and IL-2 protein production from sorted CR2^+^ versus CR2^−^ naïve CD4^+^ T cells after activation and verified the RNA results showing that IL-8 is preferentially produced by the CR2^+^ subset whereas the opposite is the case for IL-2 (**Figure 4C**). The measurement of percent positive for IL-8 is an underestimate of the difference between CR2^+^ versus CR2^−^ naïve CD4^+^ T cells since for the CR2^+^ cells expressing IL-8 the production of IL-8 on a per cell basis was greater than for CR2^−^ cells. The opposite was observed for IL-2: the few CR2^+^ cells producing IL-2 produced less on a per cell basis than CR2^−^ cells. Because CR2 rapidly disappears from the surface of T cells activated *in vitro*, it prevents the analysis of cytokine production using this marker after stimulation unless cells are sorted by CR2 expression first as in **Figure 4C**. Therefore, we correlated IL-8 production in CD4^+^ T cells defined by CD45RA, CD31 and CD25 expression with the frequency of CR2^+^ cells within the CD31^+^ CD25^−^ naïve T cell subset in samples from cord blood, MS patients reconstituting their naïve T cell compartment and MS patients in whom naïve T cells had undergone over a decade of homeostatic expansion (**Figure 4D**). Notably all MS patients that were less than one year post-depletion (n=4) had CR2 expression on 70% or more of their naïve CD4^+^ T cells (similar to the patients described in **Figures 3, S3**). Approximately 50% of these naïve CD4^+^ T cells produced IL-8; IL-8 production by naïve cells from these patients was as prevalent as that observed from cord blood naïve CD4^+^ T cells (n=3). These data support the hypothesis that RTEs in adults recapitulate the developmental stage observed in neonatal RTEs. On the other hand, the frequencies of naïve CD4^+^ T cells expressing CR2 and CD45RA^+^ CD4^+^ T cells producing IL-8 in MS patients >10 years after depletion (n=3) was less than 35% and 12%, respectively. A correlation between CR2 expression and IL-8 production was therefore observed in the seven MS patients examined. Where homeostatic expansion had occurred for at least a decade (**Figure 4D**) MS patients were similar to heathy adults (**Figure 4C**): fewer of their naïve CD4^+^ T cells are CR2+ and a lower percentage of the CR2^+^ cells produce IL-8 (0-34%). Since the expression of CR2 on a per cell basis is lower in adults than in neonates and children (**Figure 2A**), and the number of sjTRECs is reduced in cells with lower CR2 expression on a per cell basis (**Figure 2C**), these data indicate that as naïve cells expand homeostatically, CR2 expression decreases as well as their ability to produce IL-8.

### CR2^+^ Memory Cells Produce IL-8

When analysing IL-8 production by CD4^+^ T cells we noted that in MS patients more than ten years past lymphocyte depletion, a small fraction of activated CD45RA^−^ memory CD4^+^ T cells produced IL-8 (**Figure 4D**). We therefore hypothesised that CD4^+^ memory T cells can be expanded from IL-8 producing CR2^+^ naïve T cells *in vivo*. CR2 expression was observed on a proportion of central and effector memory CD4^+^ T cells and Tregs expressed the lowest levels of CR2 (**Figure 5A, Figure S5A**). CR2 expression on memory cells correlated with CR2 expression by naïve T cells (**Figure 5B**) and was age dependent (**Figure 5C**) suggesting that with age the CR2^+^ memory cells either lose CR2 expression or the subset contracts due to competition with CR2^−^ memory cells. CR2^+^ memory CD8^+^ T cells were also observed and their frequency was similarly age dependent (**Figure S5B**). As seen with the equivalent CD4^+^ naïve T cell subsets, RNA analysis of sorted CR2^+^ and CR2^−^ central memory CD4^+^ T cells showed CR1 to be the most differentially expressed gene (**Figure 5D, Spreadsheet S4, Table S1**), a phenotype confirmed at the protein level (**Figure 5E**). An overall expression analysis of the CR2^+^ and CR2^−^ naïve and memory cell subsets confirmed that the two memory populations clustered together away from the two naïve subsets confirming that the CR2^+^ memory cells are bona fide memory cells, not an unusual naïve cell subset (**Figure S5C**). Sorted CR2^+^ memory cells produced higher levels of IL-8 after activation as compared to CR2^−^ memory cells, and unlike naïve CR2^+^ cells (**Figure 4C**), all memory cells producing IL-8 also produced IL-2 (**Figure 5F**).

**Figure 5.**
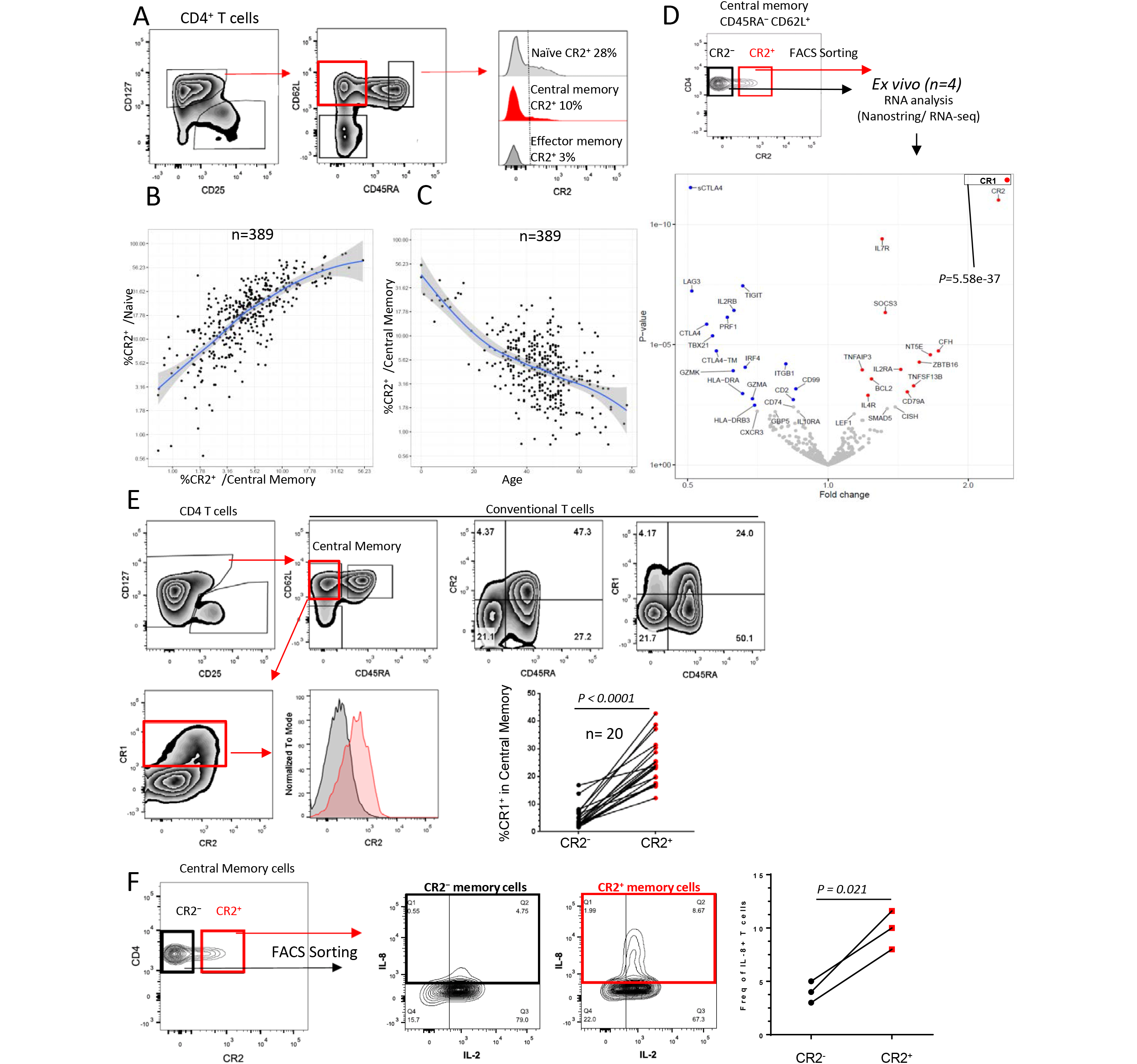
CR2 and CR1 are co-expressed on a subset of memory CD4^+^ T cells. **(A)** Gating strategy and CR2 expression on memory T cell subsets. Correlation of CR2 expression on memory versus naïve CD4^+^ T cells **(B)** and CR2^+^ memory CD4^+^ T cells versus age **(C)**. **(D)** Gating example of memory cells sorted by CR2 expression and gene expression analysis (NanoString). Color coding is described in **Figure 1. (E)** FACS analysis of CR1 and CR2 co-expression (age range 0 to 17). **(F)** Example and compiled data of IL-8 production from sorted and activated CR2^+^ and CR2^−^ memory CD4^+^ T cells (n=3, age range 30-45).

Amongst the differentially expressed genes between the CR2^−^ and CR2^+^ memory cells, a gene of particular note is *complement factor H (CFH)*, which is upregulated in both CR2^−^ and CR2^+^ memory cells as compared to naïve cells but has 2.7-fold higher levels in CR2^+^ memory cells compared to CR2" memory cells (**Figure 5D, Spreadsheet S4, Table S1**). Genes with shared expression differences between CR2^+^ naïve and CR2^+^ memory T cells versus their CR2^−^ counterparts *ex vivo* are highlighted in **Figure 6** and include a potentially relevant transcription factor for CR2^+^ CD4^+^ cells, *ZNF462*, which is expressed 13.0-fold higher in CR2^+^ versus CR2^−^ naïve cells and 7.2-fold higher on CR2+ versus CR2^−^ memory cells. Four genes, *ADAM23, ARHGAP32, DST* and *PLXNA4*, shared by the two CR2+ subsets may enhance migratory properties that augment host surveillance (28).

**Figure 6.**
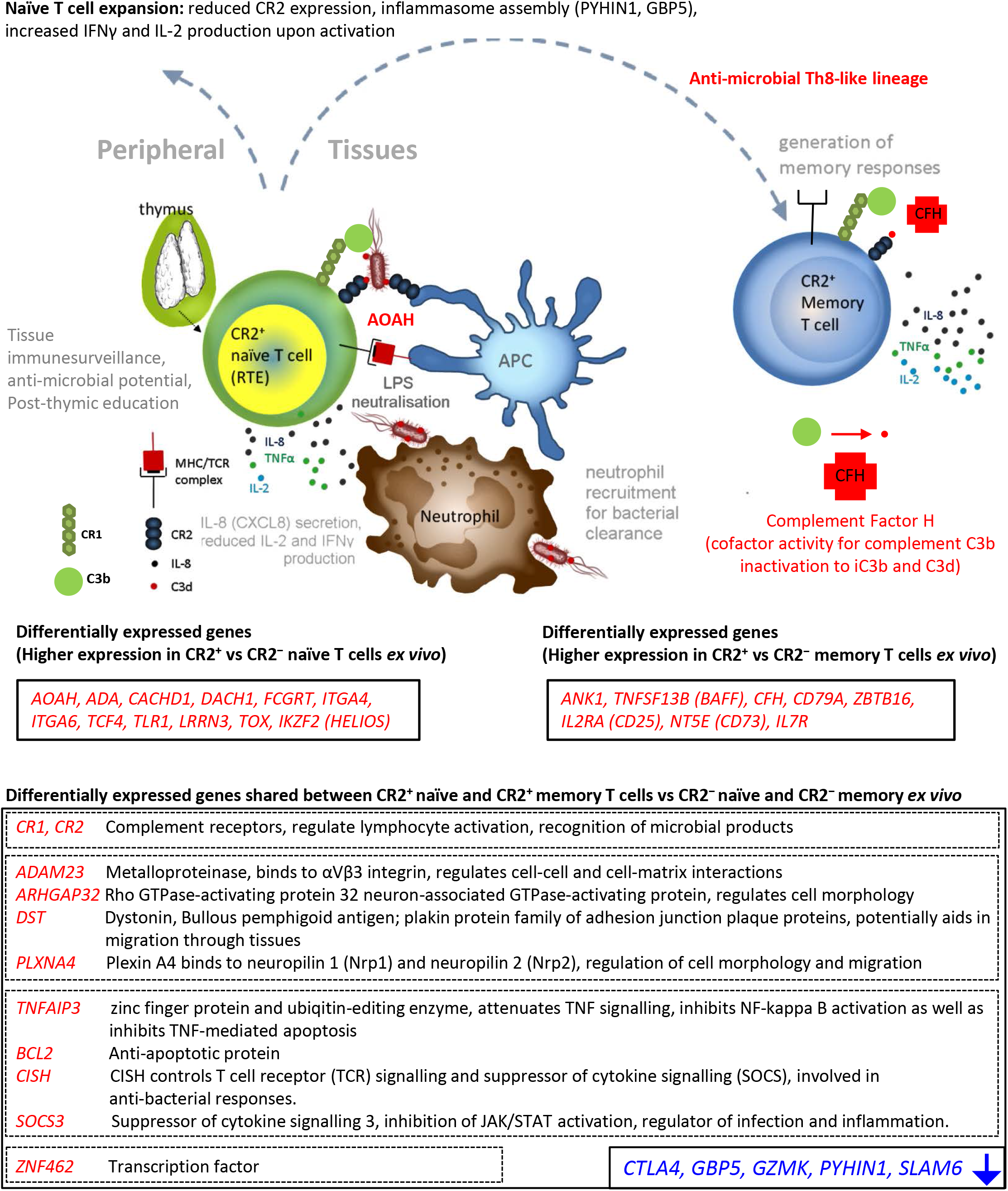
Gene expression in CR2^+^ versus CR2^−^ T cells *ex vivo* and following activation. Gene names shown in the first two boxes are some of the differentially expressed genes having higher expression in either CR2^+^ naïve or CR2^+^ memory CD4^+^ T cells versus their CR2" counterparts. Genes that share their expression patterns in both CR2^+^ naïve and memory cells are listed in the larger boxes (genes up-regulated in red, genes down-regulated in blue).

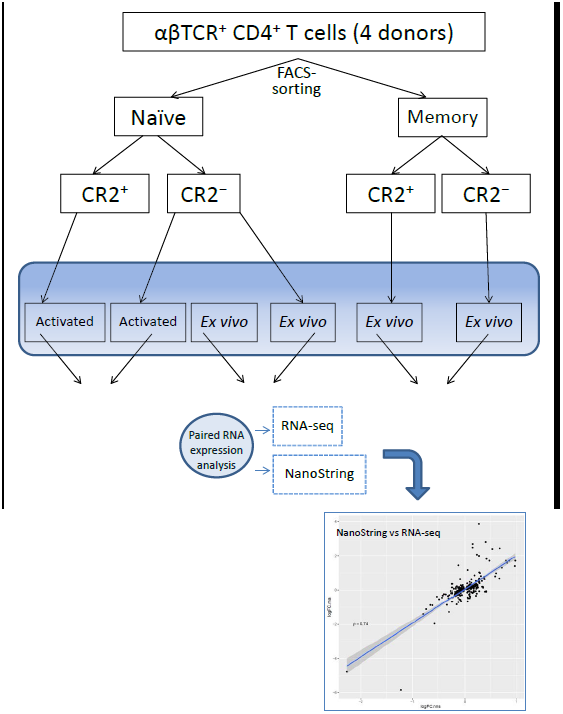
**Design of the RNA analysis experiment performed on naïve CR2^+^ and CR2^−^and memory CR2^+^ and CR2^−^ CD4^+^ T cells.** Two RNA analysis methods, RNA-seq and NanoString (Human Immunology gene panel), were applied on paired RNA samples purified from FACS-sorted T-cell subsets from four donors. Fold change (Log2) differences obtained from RNA-seq and NanoString were highly correlated (example shown at the bottom right represents fold change between CR2+ and CR2^−^ memory CD4^+^ T cells on both platforms).

## DISCUSSION

An understanding of the impact of thymic involution on health remains incomplete despite its influence on the aging of the immune system (1) and other clinical areas (11-14). In our analysis of naïve T cell subsets we report for the first time co-expression of CR1 and CR2 on the most naïve T cells and that these, as well as other molecules differentially expressed by these cells, will enable a more complete understanding of human RTEs.

Co-expression of CR1 and CR2 occurs on follicular dendritic cells and B cells and their functions are critical to generating antibody responses, including the retention of immune complexes on dendritic cells (29, 30). CR1 participates in the degradation of activated C3 to C3d, which then binds CR2 and facilitates the interaction via immune complexes of follicular dendritic cells with B cells, lowering their threshold of activation. The presence of CR2 and CR1 on the surface of naïve T cells provides them with the potential to participate in immune responses in a manner analogous to follicular DCs and B cells. Our findings add to an increasing appreciation of the role in T cell functions of molecules classically described as belonging to the innate immune system. Previously, it has been shown that human T cells express and process C3 to C3a and C3b and binding of C3b by CD46 regulates cytokine production and effector differentiation (15, 31). Although not put into the biological context demonstrated in the current study, there have been previous reports of CR1 and CR2 expression on fetal T cells (32) and thymocytes (33). A demonstration of CR2 and CR1 function in T cells was their facilitation of C3-dependent HIV infection in a cell line (34).

In addition to the complement receptors CR1 and CR2, CR2^+^ CD31^+^CD25^−^ RTEs were shown to have increased expression of the genes encoding TLR1 bacterial pattern recognition receptor, as well as AOAH, a secreted enzyme that inactivates LPS, suggesting that RTEs have a unique potential to respond as compared to long-term peripheral naïve T cells. This was further supported by our finding that following activation CR2^+^ CD31^+^CD25^−^ RTEs preferentially produced IL-8 (CXCL8), characterized as a "proinflammatory immunoprotective cytokine of neonatal T cells" via neutrophil recruitment and co-stimulation of γδ T cells (27). These authors also demonstrated that IL-8 expression by neonatal T cells was also increased when in addition to TCR stimulation cells were provided with bacterial flaggelin or the TLR1/2 agonist Pam3Cys. In our study, not only have we observed that approximately 50% of naïve CD4^+^ T cells in cord blood produce IL-8 upon activation but we also demonstrated a similar proportion of IL-8-producing naïve T cells in the blood of MS patients with newly-generated naïve T cells appearing following depletion with alemtuzmab (**Figures 3, S3**). In heathy adults or in MS patients greater than 10 years following lymphoctye depletion, a smaller portion of CD2^+^ CD31^+^ naïve cells produced IL-8, but we also noted that the level of CR2 on a per cell basis is much lower on adult CR2^+^ T cells as compared to the level of CR2 expression observed in cord blood, blood from children as well as adults actively reconstituting their immune system. This is consistent with the observation that in adults naïve CD4^+^ T cells sorted by CR2 levels the greatest number of sjTRECs were in cells with the highest CR2 levels per cell. Overall we conclude that as naïve cells emigrate from the thymus they express high levels of CR2 and are capable of secreting IL-8 but as homeostatic expansion of naïve T cells occurs through time, CR2 levels and the preferential secretion of IL-8 declines. Our findings are compatible with recent studies showing that following neonatal thymectomy IL-8 production by naïve CD4^+^ T cells is reduced by greater than 90% (8). However, with time, some children had evidence of thymic tissue regeneration that was accompanied by a higher proportion of naïve T cells secreting IL-8 following activation. We also note that a comparison of gene expression of CD31^+^ and CD31^−^ naïve CD4^+^ T cells isolated from three children between 1 and 5 years of age was made in this same study but CR2 was not observed as a differentially expressed gene, even though there were concordant results between the two studies with van den Broek and colleagues (8) reporting differential expression of genes such as *IKZF2, GBP5, PYHIN1, LRRN3, TOX* and *MAF*. In our study we noted that CD31^−^ naïve CD4^+^ T cells in neonates and children express more CR2 than the equivalent population in adults (**Figure 2A**) therefore likely accounting for the differing results for *CR2* mRNA expression between the two studies. Our results along with those of others (8) demonstrate that relatively recent development in the thymus, not absolute age, confers a unique innate phenotype to RTEs: preferential IL-8 production and high CR2 expression.

In the initial period following their emigration from the thymus T cells express high levels of CR2 and CR1, secrete IL-8 and TNF, have the capacity to hydrolyze LPS using the AOAH enzyme and produce reduced levels of T cell cytokines such as IL-2 and IFN-γ. We suggest that this differentiation state provides tissue protection while avoiding overwhelming T cell activation, attributes likely to be critical to the newborn encountering a wide range of microbes on the skin and mucosal tissues as well as encountering airborne and food antigens. This is consistent with mouse and human studies demonstrating decreased IL-2 and INF-γ production by activated RTEs compared to T cells resident in the periphery for a longer period of time (8, 35, 36). Studies focused on the biology of RTEs in mice demonstrated that in the absence of inflammation RTEs display heightened susceptibility to tolerance induction to tissue-restricted antigens (35) suggesting that the RTE differentiation state could also contribute to tolerance induction to commensals. Such tolerance induction may require RTE migration into all tissues and associated draining lymph nodes that are exposed to commensals as well as pathogens. The importance of tissue residency by naïve T cells is strongly supported by recent discoveries showing that pediatric samples of colon, ileum, jejunum and lung, in contrast to tissues from young adults, contain a large portion of naïve CD4^+^ and CD8^+^ T cells, most of which were defined as RTEs by the virtue of being CD31^+^ (36). Recently, results from TCR repertoire analyses of naïve T cells isolated from lymphoid tissues of donors aged 2 months to 73 years suggested naïve T cell homeostatic expansion is at least in part site-specific and that the dynamics of naïve T cell recirculation in humans may differ from those understood from mouse studies (4). We have found that CR2^+^ T cells express a selection of genes that give them the potential to migrate to tissue, and in the presence of appropriate danger signals and tissue-based antigen presenting cells, respond to pathogens by mediating IL-8-dependent neutrophil recruitment and differentiating into memory cells, some of which maintain the ability to secrete IL-8. This scenario of priming within tissues is supported by the presence in children of a higher proportion (>35%) of effector memory CD4^+^ and CD8^+^ T cells in jejunum, ileum and lung tissue samples as compared to lymph nodes (<10% Teff) (36). The hypothesis that RTEs seed and survey tissues throughout the body is supported by the preferential expression of a number of molecules associated with cell homing and movement by CR2+ as compared to CR2^−^ CD31^+^ cells including the gut-homing integrin alpha4 *(ITGA4)(37)* and integrin alpha6 (*ITGA6*, CD49f), a hemidesmosomal component important for tissue homing (38, 39). CR2^+^ naïve and memory T cells share the expression of other genes that are involved in cell adhesion and may enhance the migratory properties of the T cells through the basal lamina: *DST* that encodes an adhesion junction protein anchoring intermediate filaments to hemidesmosomes(40, 41), *ARHGAP32 (RICS)* that regulates neuronal cell morphology and movement(42), and *PLXNA4* that is also involved in nerve fiber guidance(43).

Direct comparison of CR2^+^ RTEs to CR2^−^ CD31^+^CD25^−^ provided insight into the earliest changes that occurred when RTE undergo initial peripheral proliferation. Consistent with the recent discovery that inflammasomes are required for Th1 T cell immunity and INF-γ production(16), we observed that increased production of IL-2 and IFN-γ in homeostatically expanded naïve T cells was correlated with the expression of *GBP5* and *PYHIN1*, molecules involved in assembly of functional inflammasomes and upregulated in CD31^−^ naïve T cells. This also suggests that as we age naïve T cells make use of differing pathogen sensor mechanisms including complement and inflammasomes, and that naïve T cells may start relevant "Th" differentiation programs during their post-thymic maturation and homeostatic expansion. Therefore, naïve T cells at different stages of post-thymic development may form functionally different memory responses. The high frequency of CR2^+^ memory T cells in children is consistent with this hypothesis.

Aspects of the innate signature of RTEs are retained by a subset of CR2^+^ memory T cells that express CR1 and secrete IL-8 upon activation, suggesting there is selection because of these functional attributes, consistent with the hypothesis that RTE-specific gene expression confers a functional competence retained by particular memory T cells possibly because of their complement-dependent reactivity to pathogens (see genes shared by CR2^+^ naïve and memory cells **Figure 6**). Further supporting this hypothesis we noted that when compared to their CR2^−^ counterparts, CR2^+^ memory T cells express 3-fold higher *CFH*, a complement regulatory glycoprotein possessing a cofactor activity for complement inactivation on the surface of the host cells, but not on that of the pathogen(20). This suggests that CR2^+^ memory T cells, many of which secrete IL-8, belong to specialized anti-microbial subsets participating in complement C3-dependent pathogen clearance. However, since we did not observe differential expression of the genes encoding Th-defining transcription factors(44) *RORC* (Th17), *GATA3* (Th2) *PRDM1* (Tfh) and *BCL6* (Tfh) (although differential expression of *TBX21* (Th1) was observed, **Spreadsheet S4**) when comparing CR2^+^ and CR2^−^ memory cells, this implies that CR2+ naïve T cells can differentiate to Th memory lineages not characterized by IL-8 secretion while maintaining CR2 expression. Genome-wide expression analysis at a single-cell level is required to clarify the heterogeneity of CR2^+^ memory cells in blood and tissues. Similar to RTEs, CR2^+^ memory cells have increased expression of genes conferring tissue migratory potential suggesting that at least a portion of this subset we have defined are a recirculating population of the recently reported IL-8^+^-producing memory CD4^+^ and CD8^+^ T cells identified within the adult skin and other tissues(17). Wong and colleagues noted the possible developmental relationship of IL-8-producing memory cells in tissues with the abundant IL-8-secreting naïve T cells in cord blood and noted that "while not defined as a Th lineage, IL-8-producing cells had very little overlap with other Th subsets in terms of cytokine secretion and trafficking receptor expression." We suggest that the shared gene expression pattern of memory and naïve CR2^+^ cells that preferentially secrete IL-8, including the transcription factor ZNF462, support a developmental relationship.

Our findings further challenge the traditional view that particular innate, germline-encoded mechanisms to identify pathogens and metabolic hazards are exclusive to the innate immune system. The majority of CR2^+^ T cells emigrate from the thymus during the childhood and their potential to interact with microbes is likely involved in priming of the T cell compartment to pathogens as well as providing mechanisms to allow tolerance to commensals. The presence of EBV receptors (CR2 molecules) on T cells highlights a potential pathway of EBV infection that in some cases results in T cell lymphoma(45). In addition, it is possible that that the binding of EBV via CR2 to EBV-specific, CR2^+^ naïve T cells during antigen-specific activation could modulate responsiveness thereby accounting for the observation that the severity of EBV increases with age(46), which correlates with the loss of CR2 on naïve T cells with homeostatic expansion. The near absence of CR2^+^ T cells later in life could contribute to the reduced immunity, especially to microbial infections, observed in older individuals(3).

## EXPERIMENTAL PROCEDURES

### Human samples

Donors of peripheral blood volunteered for one of four studies that are detailed in the Supplemental Methods.

### Whole blood and PBMC immunostaining

Blood samples were directly immunophenotyped within 5 hours following donation. Samples were blocked for 10 min with mouse IgG (20 μg/ml), stained for 40 min at room temperature with appropriate antibodies and then lysed with freshly prepared 1X BD FACS Lysing Solution (BD Biosciences). After lysis of red blood cells, samples were washed with BD CellWASH (BD Biosciences). Finally, the samples were fixed with freshly prepared 1X BD CellFIX (BD Biosciences). The samples were stored at 4 °C in the dark until analysis using a BD Fortessa flow cytometer. PBMC samples, prepared as previously described (47), were blocked for 10 min, stained for 1 hour at 4°C, washed twice and fixed as described for peripheral blood immunophenotyping except for intracellular staining when surface-stained cells after the wash-step were placed in FOXP3 Fix/perm buffer (eBioscience). Phenotyping panels are detailed in **Supplemental Experimental Procedures**. CD25 detection sensitivity was increased (47) by simultaneous application of two anti-CD25 monoclonal antibodies labelled with the same fluorochrome (clones 2A3 and M-A251, BD Biosciences). Antibody concentrations used were based on the manufacturer's instructions as well as on optimization studies. Appropriate isotype controls and fluorescence-minus-one conditions were used during the development of staining panels. Immunostained samples were analysed on a BD LSRFortessa cell analyzer and data were visualized using Flowjo (TreeStar).

### Cryopreserved PBMC

PBMC isolation, cryopreservation and thawing were performed as previously described (47) and details are in the Supplemental Methods.

### T cell subset purification by cell sorting and DNA isolation

CD4^+^ T cells (RosetteSep Human CD4^+^ T Cell Enrichment Cocktail, STEMCELL Technologies) were washed and immediately incubated with antibodies to surface molecules (**Supplemental Experimental Procedures**) for 40 minutes at 4 °C, washed and followed by sorting on a FACSAria Fusion flow cytometer cell sorter) into X-VIVO medium (Lonza) containing 5% human AB serum (Sigma). In order to isolate DNA, sorted cell subsets were checked for purity and DNA was isolated using a DNA extraction reagent (QIAGEN).

### sjTREC assay

The sjTREC assay was performed as described previously (9) and details are in the **Supplemental Experimental Procedures**.

### T cell activation

FACS-purified T cell subsets or total CD4^+^ T cells (RosetteSep) for cord blood samples and samples from MS patients in **Figure 4D** were stimulated with either anti-CD3/CD28 beads (Life Technologies) at 3 cells per bead overnight (for RNA expression analyses) or cell stimulation reagent (PMA and Ionomycin, eBioscience) in the presence of protein transport inhibitors (eBioscience) for six hours at 37 °C in 96-well U-bottom plates (for intracellular cytokine determinations). IL-8^+^ and IL-2^+^ T cells were identified with a staining panel shown in **Supplemental Experimental Procedures**.

### Microarray gene expression analysis

Total RNA was prepared from cell subsets isolated by sorting using TRIzol reagent (Life Technologies). Single-stranded cDNA was synthesised from 200 ng of total RNA using the Ambion WT Expression kit (Ambion) according to the manufacturer's instructions. Labelled cDNA (GeneChip Terminal Labelling and Hybridization Kit, Affymetrix) was hybridized to a 96 Titan Affymetrix Human Gene 1.1 ST array.

Power calculations to determine the sample size required were performed using the method of Tibshirani (48), using a reference dataset from the Affymetrix GeneST array (deposited in ArrayExpress (http://www.ebi.ac.uk/arrayexpress/, accession number E-MTAB-4852) using the TibsPower package [http://github.com/chr1swallace/TibsPower] in R. We chose 20 pairs to have a false discovery rate (FDR) close to zero whilst detecting a 5-fold change in gene expression in 20 genes with a false negative rate of 5% or a 2-fold change in gene expression in 20 genes with a false negative rate of 40%, at a significance threshold of 10^-6^.

Microarray gene expression log2 intensities were normalised using vsn2 (49). Analysis of differential expression (log2 intensities) was conducted pairwise between each cell subtype using paired t tests with limma (50). P values were adjusted using the Benjamini-Hochberg algorithm. Illustrative principal component analysis was performed on the union of the most differentially expressed genes in each pairwise comparison, the list of which is available as a Table S1E. Data are deposited with ArrayExpress, accession number E-MTAB-4853.

### NanoString and RNA-seq: sample preparation and data analysis

See **Supplemental Experimental Procedures** for a description of the NanoString and RNA-seq experimental design. CR2^+^ and CR2^−^ naïve and memory cell subsets isolated by sorting CD4^+^ T cells (RosetteSep) from four donors were pelleted directly or following activation and lysed in QIAGEN RLT buffer and frozen at -80 °C. To extract RNA, lysates were warmed to room temperature and vortexed. The RNA was extracted using Zymo Research Quick-RNA MicroPrep kit following the manufacturer's recommended protocol including on-column DNA digestion. RNA was eluted in 6 μl of RNase-free water. NanoString RNA expression analysis was performed using the Human Immunology v2 XT kit, 5 μl of RNA (5 ng/μl) was used per hybridisation and set up following the recommended XT protocol. Hybridisation times for all samples were between 16 and 20 hours. A NanoString Flex instrument was used and the Prep Station was run in high sensitivity mode and 555 fields of view were collected by the Digital Analyser. For RNA-seq analysis, 10 μl of RNA (8 ng/μl) were processed by AROS Applied Biotechnology using the Illumina TruSeq Access method that captures the coding transcriptome after library prep.

Raw NanoString expression measurements were normalized with application of NanoString software (nSolver 2.5). Subsequently, a paired differential expression analysis was carried out using DESeq2 v1.12.3(51), with preset size factors equal to 1 for all samples. Analyses were performed using a FDR of 0.05%. A missing FDR is reported for genes that were found to contain an expression outlier by DESeq2 Cook's distance-based flagging of p-values. NanoString data are deposited with ArrayExpress, accession number E-MTAB-4854.

RNA sequencing yielded on average 35.9 million paired-end reads per library. Maximum likelihood transcript read count estimates for each sample were obtained with Kallisto v0.42.5 (52), using Ensembl Release 82 (53) as a reference transcriptome. Gene expression estimates were derived by aggregating all their constituent transcript read counts, which were then employed to perform a paired differential expression analysis using limma v3.28.5 (50). Analyses were performed using a FDR of 0.05%. A missing FDR is reported for genes that did not contain at least 2 counts per million (CPM) in at least 2 samples. Data from RNA-seq are deposited with the European Nucleotide Archive, http://www.ebi.ac.uk/ena, accession number EGAS00001001870.

### Statistical analysis of flow cytometry data interrogating T-cell subsets

Statistical analyses of a flow cytometry data were performed and presented using Prism 5 software (Graphpad.com) unless otherwise stated. Comparisons between cell subsets were performed using a paired t test unless otherwise stated. *P* < 0.05 was considered significant, error bars show the SD of the samples at each test condition.

A non-parametric method, LOESS (54), performed with R software (http://www.R-project.org) was used to analyze some of the datasets The grey zones define a 95% confidence interval for each regression line.

## SUPPLEMENTAL INFORMATION

Supplemental information includes five figures, six tables and supplemental experimental methods.

## AUTHOR CONTRIBUTIONS

MLP, JAT, LSW co-designed the study, evaluated the results and co-wrote the paper. ARG and CW contributed to the design of the study, evaluated the results and edited the manuscript. CW performed statistical analysis of microarray data. ARG performed statistical analysis of NanoString and RNA-seq data. MLP, RCF, DR, DS, MM, JB, NS, XCD, SM, SD, AJC and CS-L performed experiments. HS coordinated sample collection and processing. NW coordinated sample and data management. AJColes and JLJ co-designed and co-supervised the alemtuzumab study and edited the manuscript. DBD edited the manuscript. DBD, FW-L, LSW and JAT managed acquisition of samples from normal healthy volunteers as detailed in the Supplemental Methods section.

## ACKNOWLEDGMENTS

This research was supported by the Cambridge NIHR BRC Cell Phenotyping Hub. In particular, we wish to thank Chris Bowman and Anna Petrunkina Harrison for their advice and support in cell sorting. We thank Sarah Dawson, Pamela Clarke, Meeta Maisuria-Armer and Gillian Coleman for their help in processing blood samples and Jane Kennet and Katerina Anselmiova for coordinating and obtaining blood samples from the Investigating Genes and Phenotypes Associated with Type 1 Diabetes study. We thank Karen May for obtaining blood samples from MS patients and Thaleia Kalatha for determining sample availability from patients. We thank Emma Jones for the use of the NanoString Instrument and Sarah Howlett for editorial review of the manuscript. We gratefully acknowledge the participation of all NIHR Cambridge BioResource volunteers, and thank the NIHR Cambridge BioResource centre and staff for their contribution. We thank the National Institute for Health Research and NHS Blood and Transplant. We thank the NIHR/Wellcome Trust Clinical Research Facility. This work was funded by the JDRF (9-2011-253), the Wellcome Trust (091157) and the National Institute for Health Research (NIHR) Cambridge Biomedical Research Centre. CW was supported by a Wellcome Trust grant (089989). The research leading to these results has received funding from the European Union's 7th Framework Programme (FP7/2007-2013) under grant agreement no.241447 (NAIMIT). The study was supported by the European Union's Horizon 2020 Research and Innovation Programme under grant agreement 633964 (ImmunoAgeing).

## SUPPLEMENTAL EXPERIMENTAL PROCEDURES

### Human samples

Donors of peripheral blood volunteered for one of three observational studies: Genes and Mechanisms in Type 1 Diabetes in the Cambridge BioResource recruited via the NIHR Cambridge BioResource (N=371, 114 males, 257 females; all self-reported as healthy except for four of the female donors self-reported autoimmune disease—two with autoimmune thyroid disease, one with vitiligo and one with coeliac disease, donors) was approved by NRES Committee East of England - Norfolk (ref: 05/Q0106/20); Diabetes—Genes, Autoimmunity, and Prevention, a study of newly diagnosed children with T1D and non-diabetic siblings of children with T1D (7 males aged 5 to 16, 8 females aged 1-14; all donors who participated in the current study did not have diabetes and were negative for T1D-related autoantibodies); and Investigating Genes and Phenotypes Associated with Type 1 Diabetes (6 cord blood samples; 6 adults, none with self-reported autoimmunity). Diabetes—Genes, Autoimmunity, and Prevention was originally approved by the National Research Ethics Committee London - Hampstead and is now held under the ethics of Investigating Genes and Phenotypes Associated with Type 1 Diabetes, which was approved by NRES Committee East of England - Cambridge Central (ref: 08/H0308/153). NIHR Cambridge BioResource donors were collected with the prior approval of the National Health Service Cambridgeshire Research Ethics Committee. DBD, LSW and JAT co-designed the Diabetes—Genes, Autoimmunity, and Prevention study. FW-L helped organise the provision of the blood samples in the Investigating Genes and Phenotypes Associated with Type 1 Diabetes study. JAT led the Investigating Genes and Phenotypes Associated with Type 1 Diabetes and Genes and Mechanisms of Type 1 Diabetes in the Cambridge BioResource studies.

MS patients (6 females and 2 males, aged 27-49) were from the placebo arm (received alemtuzumab only, Keratinocyte Growth Factor was not given) of the trial "Keratinocyte Growth Factor - promoting thymic reconstitution and preventing autoimmunity after alemtuzumab (Campath-1H) treatment of multiple sclerosis" (REC reference: 12/LO/0393, EudraCT number: 2011-005606-30). Additional MS patients treated with alemtuzumab greater than 10 years before sample donation were consented to a long-term follow-up study (CAMSAFE REC 11/33/0007).

### Cryopreserved PBMC

PBMC isolation was carried out using Lympholyte (CEDARLANE). PBMCs were cryopreserved in heat-inactivated, filtered human AB serum (Sigma-Aldrich) and 10% DMSO (Hybri-MAX, Sigma-Aldrich) at a final concentration of 10 x 10^6^/ml and were stored in liquid nitrogen. Cells were thawed in a 37 °C water bath for 2 min. PBMCs were subsequently washed by adding the cells to 10 ml of cold (4 °C) X-VIVO (Lonza) containing 10% AB serum per 10 x 10^6^ cells, in a drop-wise fashion. PBMCs were then washed again with 10 ml of cold (4 °C) X-VIVO containing 1% AB serum per 10 x 10^6^ cells.

### sjTREC assay

A quantitative PCR assay was purchased from Sigma-Genosys for the signal joint TCR excision circle (sjTREC) that arises through an intermediate rearrangement in the TCRD/TCRA locus in developing TCRaß^+^ T lymphocytes. An assay for the gene encoding albumin was used to normalise the data.

sjTREC.F TCGT GAGAACGGT GAATGAAG

sjTREC.R CCATGCTGACACCTCTGGTT

sjTREC.P FAM-CACGGTGATGCATAGGCACCTGC-TAMRA

Alb.F GCTGTCATCTCTTGTGGGCTGT

Alb.R ACTCATGGGAGCTGCTGGTTC

Alb.P FAM-CCTGTCATGCCCACACAAATCTCTCC-TAMRA

For each sample, 24 ng of DNA was incubated in duplicate with both primers (700 nM), probe (150 nM) and 12.5 μl TaqMan mastermix (Applied Biosystems) and processed using the Applied Biosystems™ 7900HT Fast Real-Time PCR System. sjTREC were normalized to the albumin gene, representing cellular DNA, using the following formula: 2^(Ct(albumin) - Ct(sjTREC)).

**Figure S1.**
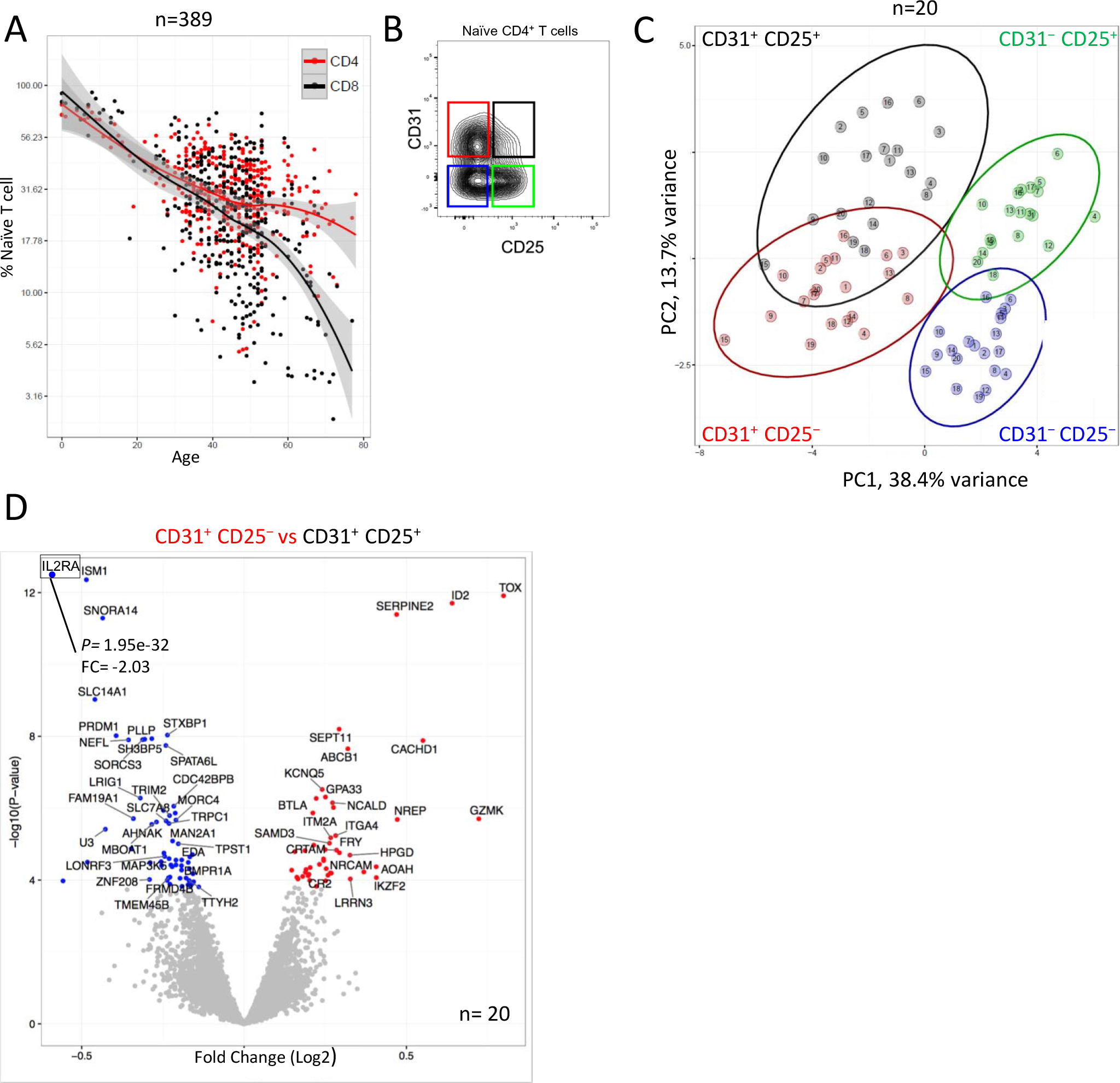
Gene expression profiling of four naïve CD4^+^ T cell subsets identify age-related molecular signatures. **(A)** Frequency of naïve CD4^+^ and CD8^+^ naïve T cells out of total CD4^+^ and CD8^+^ T cells, respectively, as a function of age (n=389). **(B)** FACS sorting strategy of naïve CD4^+^ T cells stratified by expression of CD31 and CD25. **(C)** Principal component analysis (PCA) based on differentially expressed genes amongst four naïve CD4^+^ T cell subsets stratified by surface expression of CD31 and CD25 (subsets sorted from 20 donors). **(D)** Volcano plot representing differences in gene expression between CD31^+^ CD25^−^ and CD31^+^ CD25^+^ naïve CD4^+^ T cells; genes increased in CD31^+^ CD25^−^ (red) versus those increased in CD31^+^ CD25^+^ (blue) naïve CD4^+^ T cells; *P* value (P) and Fold Change (FC) for *IL2RA* (encoding CD25) are noted since the values fall outside of the graph boundaries.

**Figure S2.**
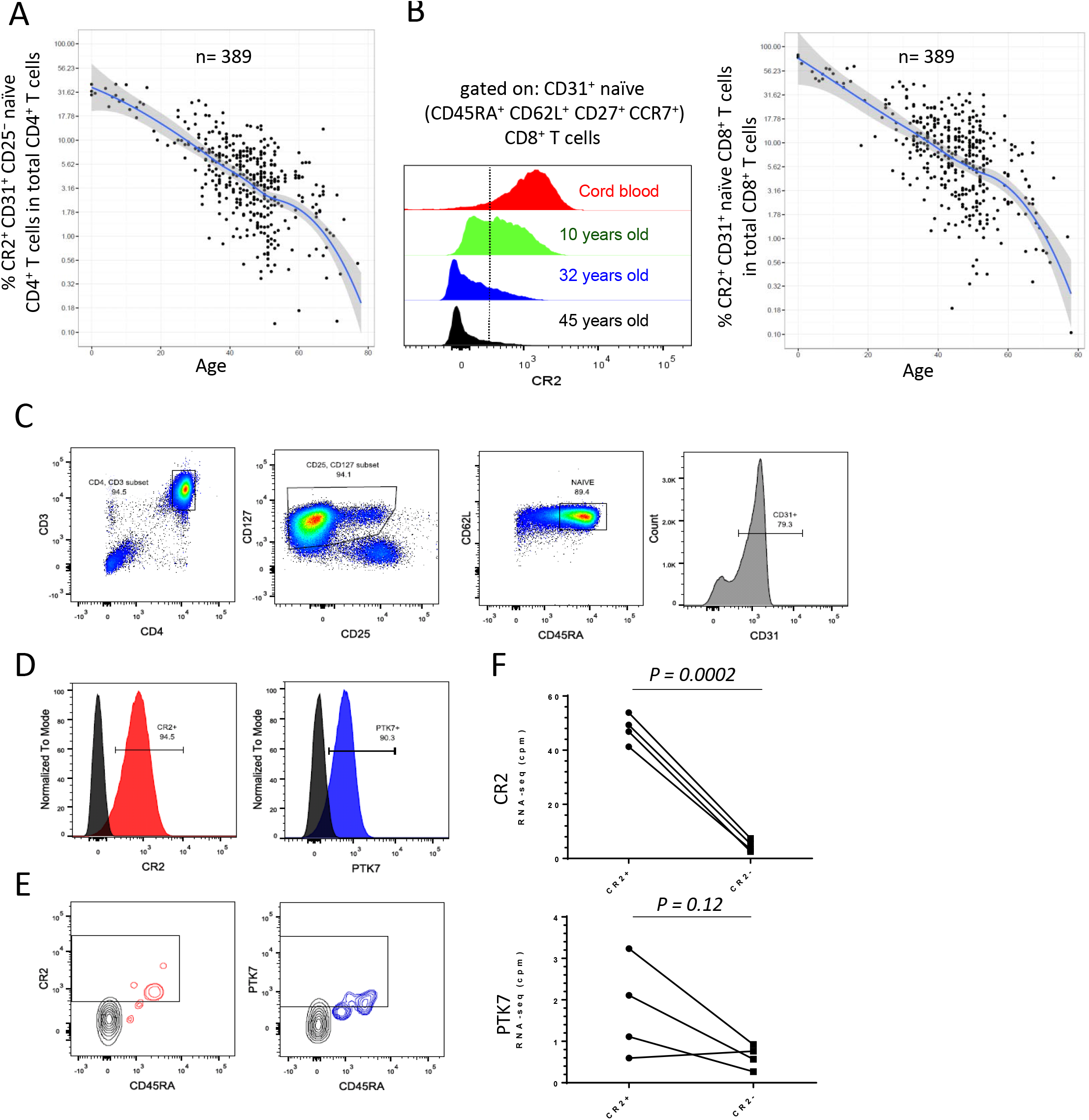
CR2 and PTK7 expression by recent thymic emigrants. **(A)** The frequency of CD31^+^ CD25^−^ naïve CD4^+^ T cells out of total CD4^+^ T cells that are CR2^+^ as a function of age. **(B)** Representative examples (from a total of 389 donors) of CR2 expression on naïve CD8^+^ T cells and the frequency of CD31^+^ naïve CD8^+^ T cells out of total CD8^+^ T cells that are CR2^+^ as a function of age (see Figure S5B for gating strategy of naïve CD8^+^ T cells). **(C)** Gating strategy of naïve CD31^+^ CD4^+^ T cells in umbilical cord blood previously enriched for CD4^+^ T cells. **(D)** Cell surface expression of CR2 and PTK7 on naïve CD4^+^ T cells from umbilical cord blood. **(E)** Representative levels of CR2 and PTK7 expression on naïve CD4^+^ T cells during T cell reconstitution (MS patient, 3 months after T-cell depletion). **(F)** Number of CR2 and PTK7 transcripts in CR2^+^ and CR2^−^ naïve CD4^+^ T cells based on RNA-seq analysis, counts per million reads (cpm); (n=4, age range 30-45, paired t test).

**Figure S3.**
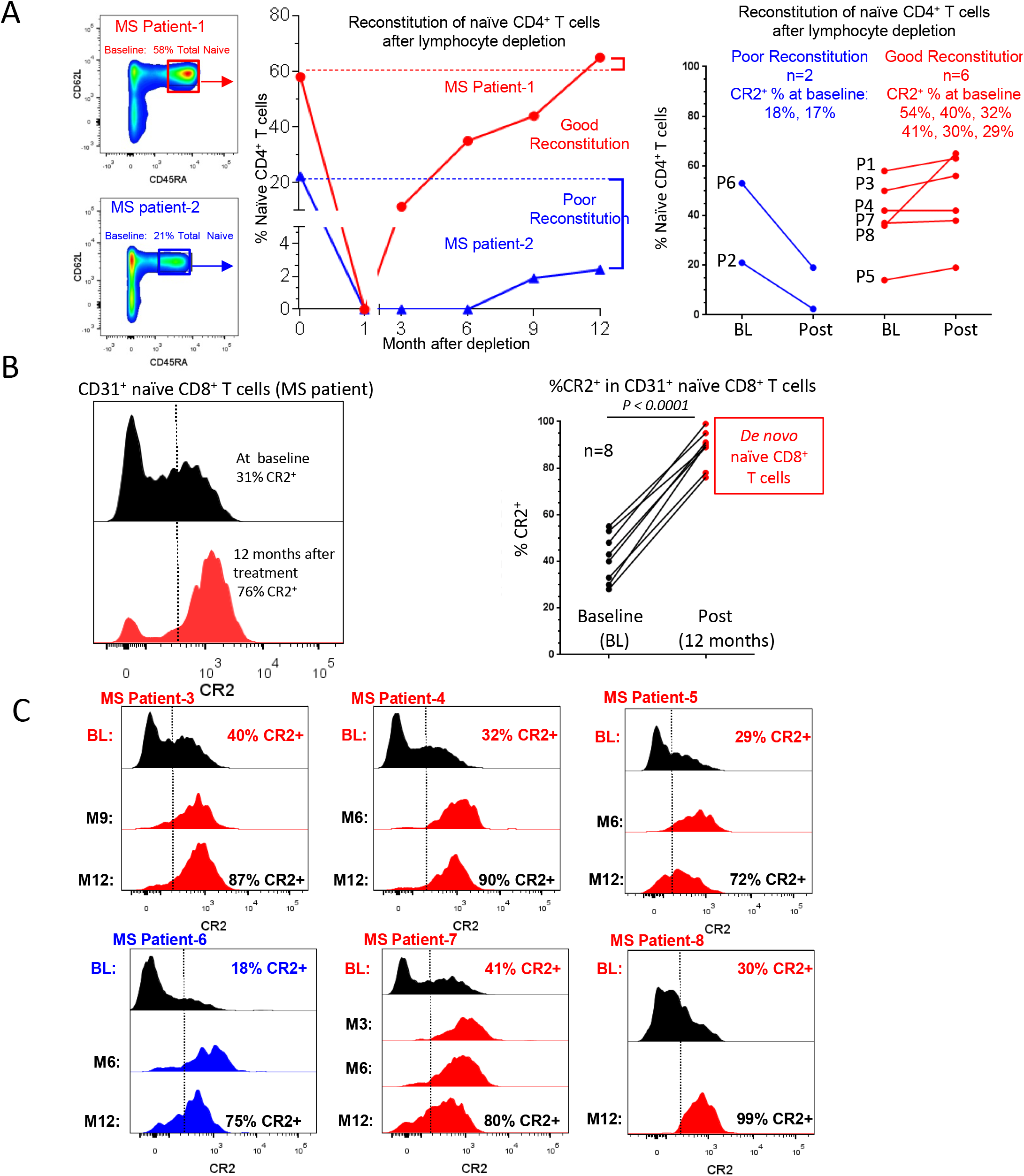
Reconstitution of naïve T cells following lymphocyte depletion in MS patients. **(A)** Two representative examples and summary of naïve CD4^+^ T cell (CD45RA^+^ CD62L^+^) reconstitution (baseline (BL) versus 12 months after depletion (Post). "P" numbers by the data points indicate the patient number. **(B)** Representative example and summary of CR2 expression on CD8^+^ naïve T cells before and 12 months after reconstitution. **(C)** CR2 expression profiles of CD31^+^ CD25^−^ naïve CD4^+^ T cells before and at time points after lymphocyte depletion (see **Figure 3** for two other patients). CR2^+^ cell frequencies in the CD31^+^ CD25^−^ naïve CD4^+^ T cell subset are shown in the baseline and month 12 histograms. Profiles depicted in red and blue denote patients with good and poor, respectively, naïve T cell reconstitution at 12 months following depletion (see **Figure S3A** above).

**Figure S4.**
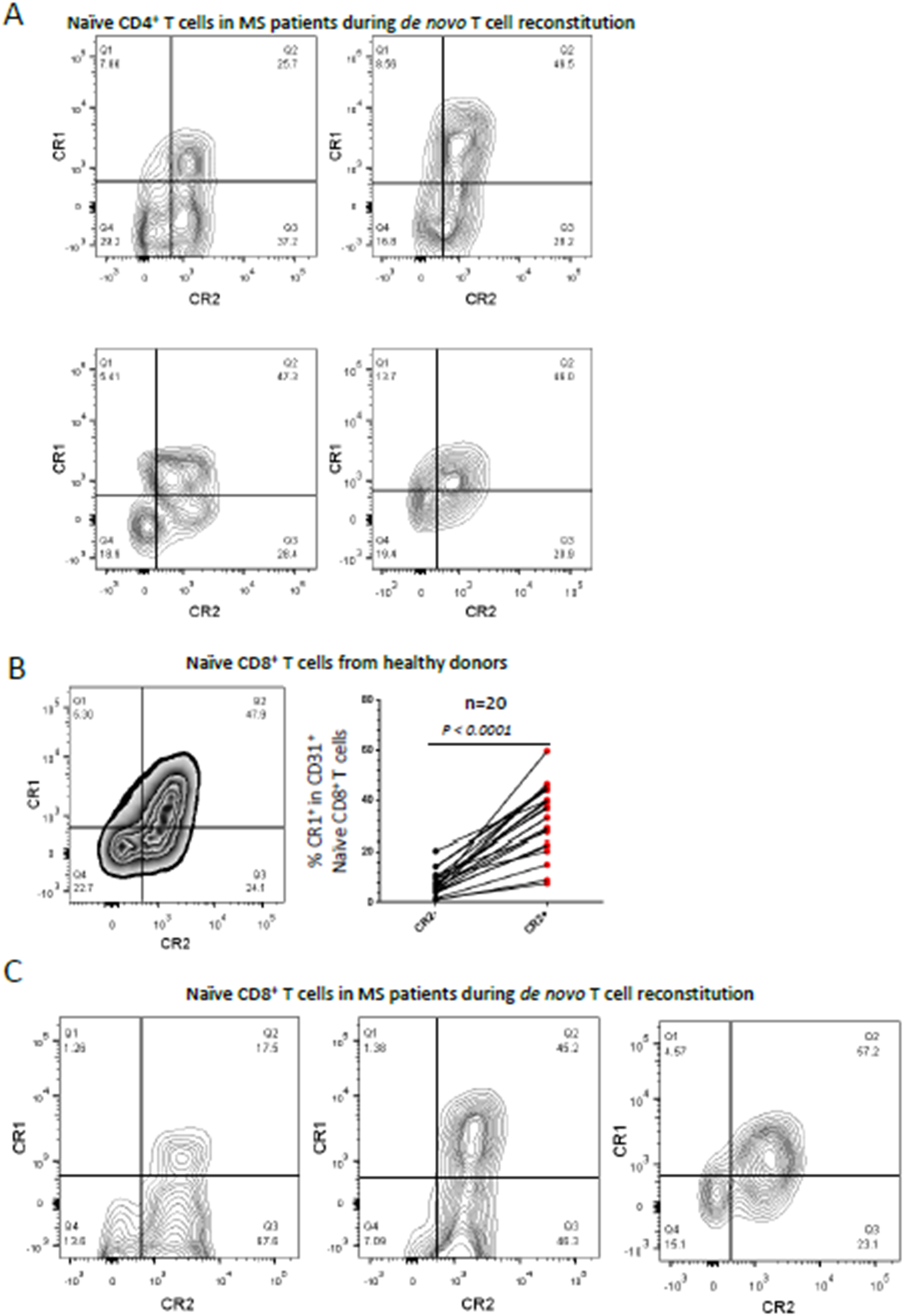
CR2 and CR1 are co-expressed on naïve T cells. **(A)** CR1 and CR2 expression on CD31^+^ CD25^−^ naïve CD4^+^ T cells present in alemtuzumab-treated MS patients during reconstitution. **(B)** Representative example and summary analysis of cell-surface CR1 expression on CR2^+^ and CR2^−^ CD31^+^ naïve CD8^+^ T cells (n=20, age range 0 to 17, paired t test). **(C)** CR2 and CR1 expression on CD31^+^ naïve CD8^+^ T cells present in alemtuzumab-treated MS patients during reconstitution.

**Figure S5.**
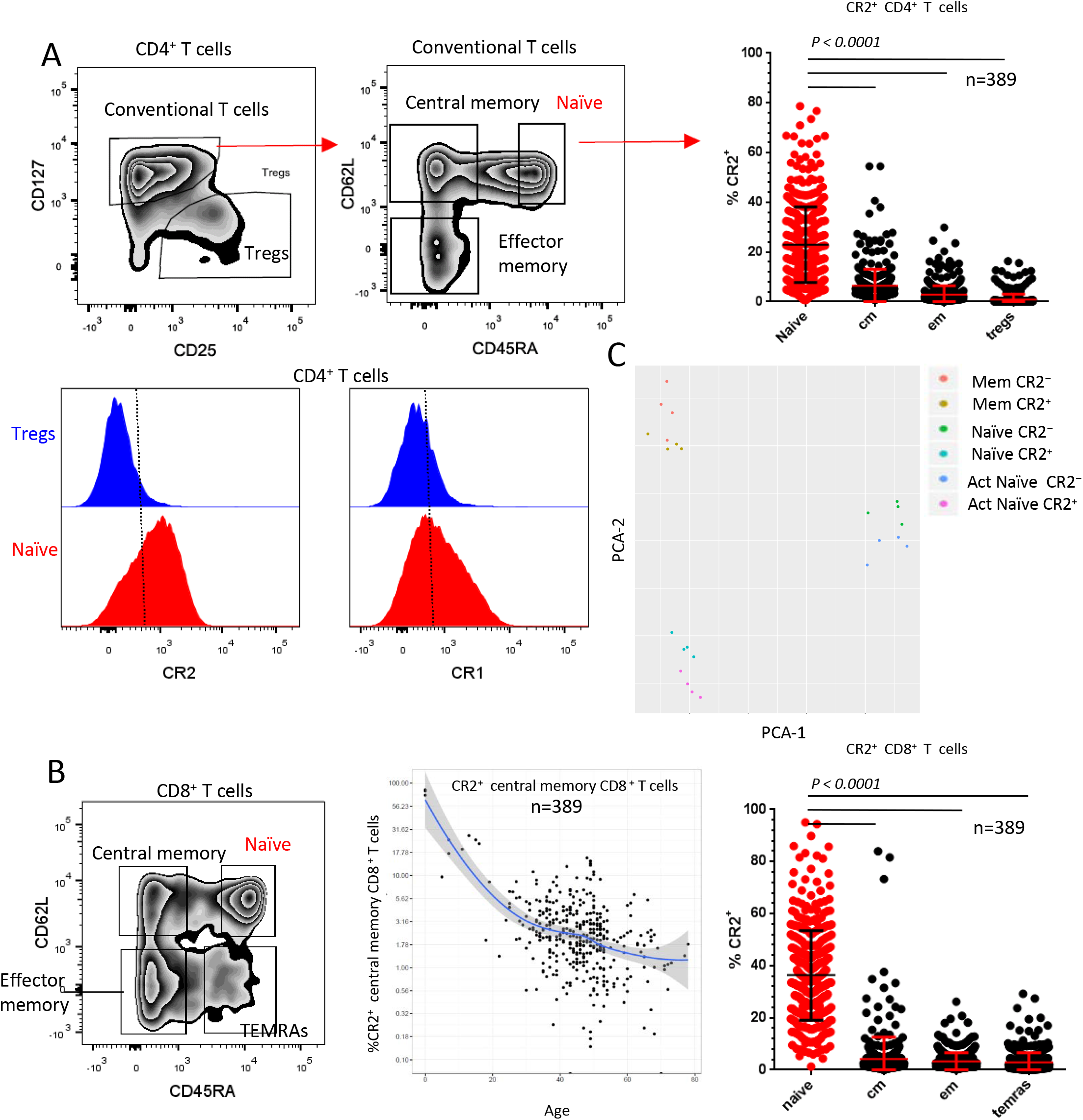
CR2 is expressed by subsets of CD4^+^ and CD8^+^ memory T cells. **(A)** Representative gating of Tregs and naïve, central memory and effector memory CD4^+^ T cells and the distribution of CR2^+^ cells within each of these subsets (paired t tests for the comparisons indicated). Representative examples of CR1 and CR2 expression on naïve CD4^+^ T cells and CD4^+^ Tregs from a donor 10 years of age. **(B)** Representative gating of naïve and memory CD8^+^ T cell subsets. CR2 expression on central memory CD8^+^ T cells as a function of age. Distribution of CR2^+^ cells within naïve and memory CD8^+^ T cell subsets (mean ± SD, paired t tests for comparisons). **(C)** Principal component analysis (PCA) based on gene expression (NanoString) of CR2^+^ and CR2^−^ naïve and memory CD4^+^ T cells *ex vivo* and activated CR2^+^ and CR2^−^ naïve CD4^+^ T cells.

**Table S1.** Selected results of RNA analysis experiment (RNA-seq) performed on CR2^+^ and CR2^−^ naïve and memory CD4^+^ T cells *ex vivo.*

|  |  | Naïve *ex vivo* |  |  | Memory *ex vivo* |  |  |
| --- | --- | --- | --- | --- | --- | --- | --- |
| Gene ID |  | CR2+ | CR2- |  | CR2+ | CR2- |  |
| **CR1** | Donor1 | 271.382* | 27.106 | FC** | 1008.486 | 39.988 | FC |
|  | Donor2 | 162.204 | 12.753 | 10.204 | 825.464 | 22.576 | 27.027 |
|  | Donor3 | 180.323 | 9.47 | Adj P value*** | 1374.156 | 37.313 | Adj P value |
|  | Donor4 | 367.677 | 56.653 | 0.001 | 619.707 | 37.049 | 0.00001 |
| **CR2** | Donor1 | 53.892 | 7.447 | FC | 53.824 | 0.881 | FC |
|  | Donor2 | 41.293 | 3.413 | 10.204 | 33.583 | 0.231 | 58.824 |
|  | Donor3 | 46.893 | 2.554 | Adj P value | 55.151 | 1.041 | Adj P value |
|  | Donor4 | 49.316 | 5.371 | 0.001 | 21.741 | 0.52 | 0.001 |
| **ZNF462** | Donor1 | 6.921 | 0.44 | FC | 31.936 | 4.507 | FC |
|  | Donor2 | 3.14 | 0.237 | 12.987 | 17.958 | 0.971 | 7.194 |
|  | Donor3 | 13.643 | 0.887 | Adj P value | 24.472 | 4.212 | Adj P value |
|  | Donor4 | 10.583 | 0.967 | 0.005 | 23.426 | 2.914 | 0.005 |
| **CACHD1** | Donor1 | 46.55 | 31.59 | FC | 1.997 | 0.102 | FC |
|  | Donor2 | 40.215 | 20.102 | 1.757 | 1.395 | 0.37 | n/a |
|  | Donor3 | 33.138 | 13.265 | Adj P value | 0.423 | 0.098 | Adj P value |
|  | Donor4 | 45.642 | 29.65 | 0.037 | 2.467 | 1.041 | n/a |
| **ADAM23** | Donor1 | 11.703 | 2.482 | FC | 94.556 | 20.536 | FC |
|  | Donor2 | 2.344 | 0.711 | 6.803 | 45.053 | 9.016 | 5.051 |
|  | Donor3 | 21.67 | 1.561 | Adj P value | 90.442 | 14.808 | Adj P value |
|  | Donor4 | 14.583 | 2.578 | 0.02 | 36.686 | 6.765 | 0.00001 |
| **PLXNA4** | Donor1 | 12.262 | 2.242 | FC | 83.232 | 20.638 | FC |
|  | Donor2 | 3.562 | 0 | 6.024 | 20.666 | 1.618 | 4.926 |
|  | Donor3 | 21.494 | 3.866 | Adj P value | 59.546 | 7.258 | Adj P value |
|  | Donor4 | 25.493 | 3.867 | 0.02 | 36.034 | 11.188 | 0.014 |
| **AOAH** | Donor1 | 19.157 | 9.009 | FC | 7.017 | 7.388 | FC |
|  | Donor2 | 64.166 | 22.709 | 2.155 | 14.26 | 9.385 | 1.185 |
|  | Donor3 | 80.594 | 25.749 | Adj P value | 12.827 | 13.116 | Adj P value |
|  | Donor4 | 129.859 | 88.895 | 0.046 | 46.509 | 32.628 | 0.512 |
| **CFH** | Donor1 | 0.037 | 0.08 | FC | 47.235 | 18.842 | FC |
|  | Donor2 | 0.094 | 0.759 | n/a | 41.539 | 23.903 | 2.653 |
|  | Donor3 | 0 | 0 | Adj P value | 45.677 | 10.805 | Adj P value |
|  | Donor4 | 0 | 0 | n/a | 58.334 | 19.254 | 0.016 |
| **DST** | Donor1 | 11.404 | 5.966 | FC | 49.574 | 20.875 | FC |
|  | Donor2 | 15.467 | 7.823 | 1.770 | 30.896 | 8.692 | 2.653 |
|  | Donor3 | 1.653 | 0.816 | Adj P value | 33.11 | 10.089 | Adj P value |
|  | Donor4 | 32.451 | 20.196 | 0.038 | 61.639 | 25.655 | 0.006 |
| **ARHGAP32**<br>**(RICS)** | Donor1 | 37.381 | 24.183 | FC | 50.943 | 19.79 | FC |
|  | Donor2 | 52.589 | 19.39 | 2.105 | 59.209 | 21.083 | 2.427 |
|  | Donor3 | 49.637 | 18.443 | Adj P value | 31.417 | 11.391 | Adj P value |
|  | Donor4 | 55.383 | 28.629 | 0.02 | 35.429 | 19.462 | 0.007 |
| **PYHIN1** | Donor1 | 5.516 | 12.772 | FC | 30.121 | 47.477 | FC |
|  | Donor2 | 8.718 | 17.589 | 0.444 | 45.466 | 68.102 | 0.703 |
|  | Donor3 | 3.694 | 9.151 | Adj P value | 43.3 | 51.65 | Adj P value |
|  | Donor4 | 3.061 | 6.07 | 0.02 | 33.66 | 44.076 | 0.039 |

\*Gene counts per million reads
\*\*FC= Fold change of CR2<sup>+</sup> vs CR2<sup>-</sup>
\*\*\*Adj P value= Adjusted P value

**Table S2.** Selected results of RNA analysis experiment (NanoString) performed on CR2^+^ and CR2∼ naïve CD4^+^ T cells after activation.

| Gene ID |  | Naïve activated |  |  |
| --- | --- | --- | --- | --- |
|  |  | CR2<sup>+</sup> | CR2<sup>-</sup> |  |
| IL8 | Donor1 | 2529* | 889 | FC** |
|  | Donor2 | 69 | 24 | 2.797 |
|  | Donor3 | 140 | 52 | Adj P value*** |
|  | Donor4 | 187 | 50 | 7.36E-21 |
| IL2 | Donor1 | 3937 | 20724 | FC |
|  | Donor2 | 9108 | 31986 | 0.307 |
|  | Donor3 | 16729 | 40604 | Adj P value |
|  | Donor4 | 11808 | 40884 | 3.52E-23 |
| IL21 | Donor1 | 3 | 56 | FC |
|  | Donor2 | 25 | 281 | 0.107 |
|  | Donor3 | 34 | 385 | Adj P value |
|  | Donor4 | 26 | 341 | 3.19E-52 |
| IFNG | Donor1 | 9 | 63 | FC |
|  | Donor2 | 8 | 17 | 0.426 |
|  | Donor3 | 14 | 36 | Adj P value |
|  | Donor4 | 36 | 84 | 3.11E-04 |
| LIF | Donor1 | 418 | 769 | FC |
|  | Donor2 | 319 | 778 | 0.489 |
|  | Donor3 | 305 | 622 | Adj P value |
|  | Donor4 | 301 | 632 | 3.70E-16 |
| TNF | Donor1 | 945 | 1130 | FC |
|  | Donor2 | 641 | 762 | 0.898 |
|  | Donor3 | 1487 | 1353 | Adj P value |
|  | Donor4 | 1580 | 1914 | 4.06E-01 |
| IL23A | Donor1 | 578 | 536 | FC |
|  | Donor2 | 248 | 248 | 1.123 |
|  | Donor3 | 463 | 317 | Adj P value |
|  | Donor4 | 441 | 430 | 4.68E-01 |
| LTA | Donor1 | 1115 | 1013 | FC |
|  | Donor2 | 1182 | 1187 | 1.137 |
|  | Donor3 | 1857 | 1362 | Adj P value |
|  | Donor4 | 2217 | 1965 | 2.66E-01 |

\*Normalized Counts
\*\*FC=Fold change of CR2<sup>+</sup> vs CR2<sup>-</sup>
\*\*\*Adj P value=Adjusted P value

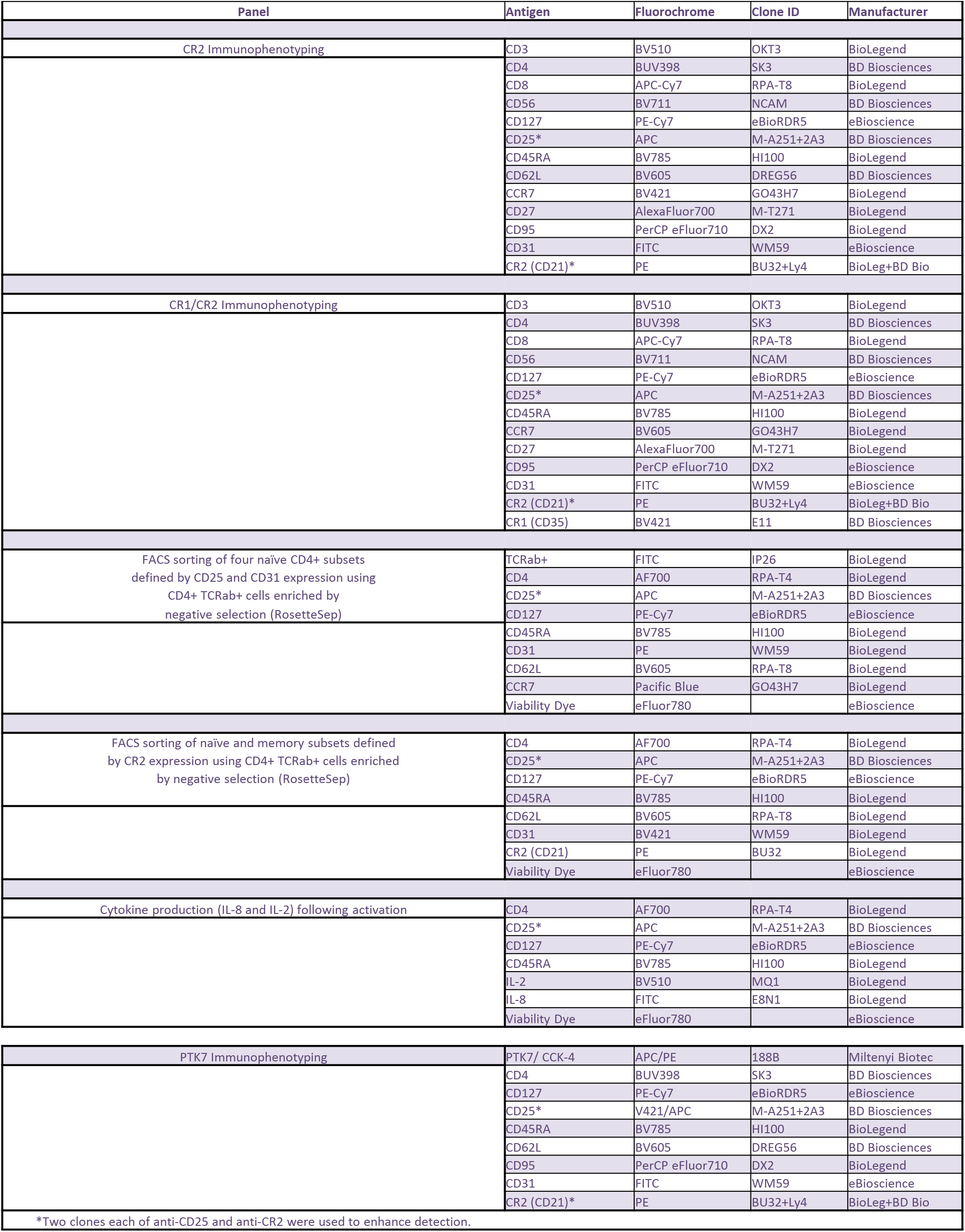
Antibody panels used for immunophenotyping and FACS-sorting.

